# Exercise modulation of the alternative splicing landscape in human tissues

**DOI:** 10.64898/2026.03.03.704460

**Authors:** Zidong Zhang, German Nudelman, Hanna Pincas, Gayatri Iyer, Gregory R. Smith, Hasmik Keshishian, Christopher A. Jin, Scott Trappe, Daniel H. Katz, Charles F. Burant, Venugopalan D. Nair, Elena Zaslavsky, Stuart C. Sealfon, the MoTrPAC Study Group

**Affiliations:** Department of Neurology, Icahn School of Medicine at Mount Sinai, New York, New York, USA; Department of Computational Medicine and Bioinformatics, University of Michigan, Ann Arbor, MI, USA; Proteomics Platform, Broad Institute of MIT and Harvard, Cambridge, Massachusetts, USA; Department of Genetics, Stanford University School of Medicine, Stanford, California, USA; Human Performance Laboratory, Ball State University, Muncie, Indiana, USA; Department of Medicine, Stanford University School of Medicine, Stanford, California, USA; Department of Internal Medicine, University of Michigan, Ann Arbor, MI, USA

**Keywords:** acute endurance exercise, acute resistance exercise, alternative splicing, epigenome, proteome, phosphoproteome, spliceosome, skeletal muscle, adipose tissue, blood

## Abstract

The diverse health benefits of exercise are associated with multi-organ molecular responses. Alternative RNA splicing (AS) is an important determinant of transcriptome and proteome diversity. We profiled the temporal effects of acute endurance and resistance exercise on the AS landscape of human skeletal muscle, adipose tissue, and blood, and studied regulatory mechanisms through integrated multi-omic analyses. We identified 5102 distinct differential AS (DAS) events, with the majority modifying protein-coding sequence (89%) and being independent of altered RNA expression (67%). Endurance and resistance exercise induced differing patterns of AS alterations with divergent temporal trajectories. We inferred the DAS-associated RNA-binding and DNA-binding proteins. In skeletal muscle, where DAS events were the most abundant, DAS genes were enriched for muscle structure- and RNA splicing-related processes, and splicing machinery components were regulated at the protein phosphorylation, RNA, and AS levels. These findings implicate AS regulation as a major mediator of the responses to exercise.

## Introduction

Exercise is associated with health benefits and a reduced risk of chronic diseases including cardiovascular disease, type 2 diabetes, obesity, cancer, and a delay in cognitive decline (for review, see ^1^). Exercise induces physiological responses in skeletal muscle (SKM) and in diverse tissues ^2^. Unraveling the molecular mechanisms involved in the response to acute exercise can help to elucidate long-term adaptations-related mechanisms. Further, this could provide the foundation for more personalized exercise recommendations. The molecular responses to acute exercise include signaling pathway alterations, changes in gene expression, and changes in epigenetic processes (for review, see ^3,4^). Various studies have characterized the transcriptomic responses to acute exercise in human SKM ^5–7^, adipose tissue ^8^, and blood ^9,10^. These transcriptomic responses vary with the exercise modality, e.g. endurance (EE) vs. resistance exercise (RE), the tissue type, and the characteristics of the study cohort including sex, age, ethnicity, hormonal status, etc. ^11^.

In addition to regulation of transcript and proteome abundance by differential transcript levels, an estimated 95% of human genes generate multiple transcripts through alternative splicing (AS; ^12^). AS differentially selects splice site(s) on the mRNA precursor, giving rise to multiple transcript variants that are referred to as splicing variants/isoforms. Thus, AS is an important co-transcriptional mechanism contributing to transcriptome complexity, RNA expression dynamics, and proteome diversity (for review, see ^13,14^). The splicing process is mediated by the spliceosome, a macromolecular complex composed of small nuclear RNAs, small nuclear ribonucleoproteins (snRNPs), and numerous proteins comprising RNA-binding proteins (RBPs), which are also referred to as splicing factors (SFs; for review, see ^15^). The regulation of AS occurs at the RNA, transcriptional, and epigenetic levels (for review, see ^16^). At the RNA level, AS is regulated by interactions between cis-regulatory elements on the mRNA precursors and SFs. SFs either promote or inhibit the use of adjacent splice sites. Additionally, AS can be modulated by core elements of the spliceosome, e.g. at the level of snRNP biogenesis, the expression of core spliceosomal proteins, or snRNA abundance (for review, see ^17^). At the transcriptional level, DNA-binding transcription factors (TFs)can lead to variation in the RNA polymerase II elongation rate to impact AS patterns (see ^18^). In addition, TFs can regulate AS by recruiting other RNA-binding SFs or by directly binding to the pre-mRNA ^19^.

AS is thought to play an important role in SKM physiology and in neuromuscular diseases (for review, see ^20^. Previous mouse studies suggest that AS may also contribute to improving athletic performance ^21–23^. However, study of its modulation by exercise has been limited, and little is known about the mechanisms underlying exercise-regulated AS. Previous studies in SKM and blood have identified the regulation of individual splicing isoforms with exercise, including hTERT, PGC-1α, VEGF, IFG-I, TNNT1, and HIF1α in human ^24–33^, and Axl, Dync1, Plekhg1, and Cobll1 in horse ^34–36^. Endurance exercise training in humans leads to regulation of a large number of transcript isoforms in SKM ^37^. A recent study including a meta-analysis of human public RNA-seq datasets from a small number of individuals reported several hundred AS events in SKM in response to acute EE ^38^. Analyzing the modulation of the AS landscape by acute EE and RE genome-wide and dissecting the regulatory layers involved in its regulation would advance our understanding of the molecular effects of exercise and of gene expression regulation.

To gain deeper insight into the molecular underpinnings of exercise-induced physiological changes in healthy individuals, the MoTrPAC Consortium has developed a comprehensive multi-omic map of the temporal response to acute exercise in SKM, adipose tissue, and blood from sedentary adults (^39^; MoTrPAC Study Group 2025 Pre-CAWG Landscape, **manuscript submitted**). Here, we analyze MoTrPAC transcriptome, chromatin accessibility, proteome, and phosphoproteome datasets obtained from human SKM, adipose tissue, and blood before and after acute EE or RE. We characterize the dynamic changes of the AS landscape in response to both exercise modalities. We identify the biological processes regulated by AS and infer the mechanisms underlying exercise regulation of AS.

## Results

### Study overview

To study the exercise-induced alterations in the AS landscape of SKM, adipose tissue, and blood and uncover regulatory mechanisms driving those changes, we utilized time-series multiome data from an acute bout of exercise (**Fig. 1A**). Healthy sedentary study participants underwent either cycling endurance exercise (EE, n=65), resistance exercise (RE, n=73) or served as non-exercise controls (n=37). While an attempt was made to match all groups by age and sex distributions, because this MoTrPAC study was suspended due to the COVID-19 pandemic, the study arms had some imbalances, especially for non-exercise control males (n=6) and females (n=31). SKM and adipose tissue biopsies were taken before exercise (Pre) and after the exercise bout; at 15 min, 3.5 hr, and 24 hr post-exercise (P15M, P3.5H, and P24H) for SKM, and at 45 min, 4 hr, and 24 hr post-exercise (P45M, P4H, and P24H) for adipose tissue. Blood samples were collected before, during (20 min [D20M] and 40 min [D40 M]), and after exercise (at 10 min [P10M], 30 min [P30M], 3.5 hr [P3.5H], and 24 hr [P24H] post-exercise) (**Fig. 1B**; for details, see Pre-CAWG Landscape, **manuscript submitted**). Tissue samples were processed for multi-omic analyses, which included deep paired read RNA-seq to improve AS resolution (average 60M paired reads), ATAC-seq, proteome, and phosphoproteome (**Fig. 1C**). Temporal changes were identified and analyzed in conjunction with pre-exercise AS levels to map the AS landscape. Integrative analyses were used to infer mechanisms responsible for exercise-regulated AS events (**Fig. 1D**).

**Figure 1.**
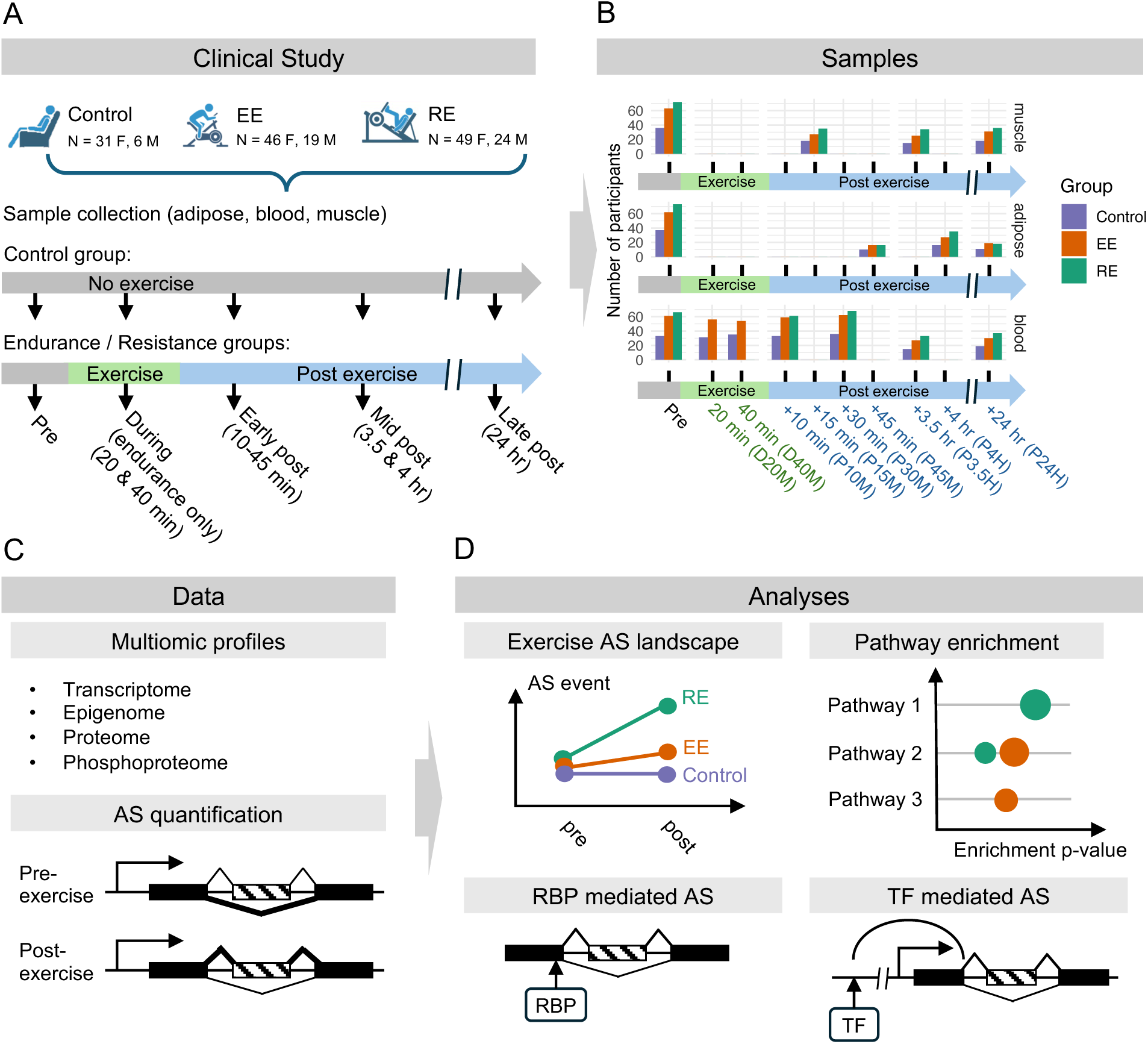
Study design, multi-omic data, and AS-related analyses. **A**, Experimental design and tissue collection. Adult sedentary participants were subjected to an acute bout of EE or RE. Participants in the control group did not perform any exercise and rested in a supine position for 40 min. Samples were collected at the indicated times. **B**, Distribution of the number of participants by group (Control, EE, and RE) across the sample collection times. **C**, Summary of the -omic datasets used in this study and of the quantification of AS events before and after exercise. Depicted is an example of a major exon skipping event detected in the Pre-exercise time point vs. leading exon inclusion in a Post-exercise time point. **D**, Summary of the analyses conducted in this study, which include studying the dynamic changes of the AS landscape, determining the biological pathways that are enriched among the genes associated with differential AS (DAS), exploring the regulation of AS by RBPs and DNA-binding TFs.

### Differential Alternative Splicing (DAS) Events

We analyzed the four most frequent types of AS events: skipped exon (SE), alternative 5’ splice site (A5SS), alternative 3’ splice site (A3SS), and retained intron (RI) (**Fig. 2A, *Left panel***). We first applied the STAR and kallisto methods to the RNA-seq fastq files to measure transcript levels with the primary processing adapted for the two AS calling methods (see Methods; ^40^). Splicing events were then determined by the isoform usage at each splice junction using a combination of two complementary statistical approaches, DARTS/rMATS-turbo ^41,42^ and SUPPA2 ^43^, as previously described ^44,45^. All analyses relied on the NCBI transcript variant annotations. The final processed data for AS analysis comprised percentage-spliced-in (PSI) values for each type of splicing event in each sample (see Methods, **Fig. S1**, and GEOxxxxxx). DAS events were identified by comparing the PSI values obtained at the pre-exercise timepoint with those at each subsequent timepoint with a logistic regression and mixed model to analyze all data while retaining within-subject comparisons (see Methods). A DAS event for either the EE or the RE group was included if it was significant either in an exercise group but not in the control group, or in both exercise and control groups with differing directions of change.

**Figure 2.**
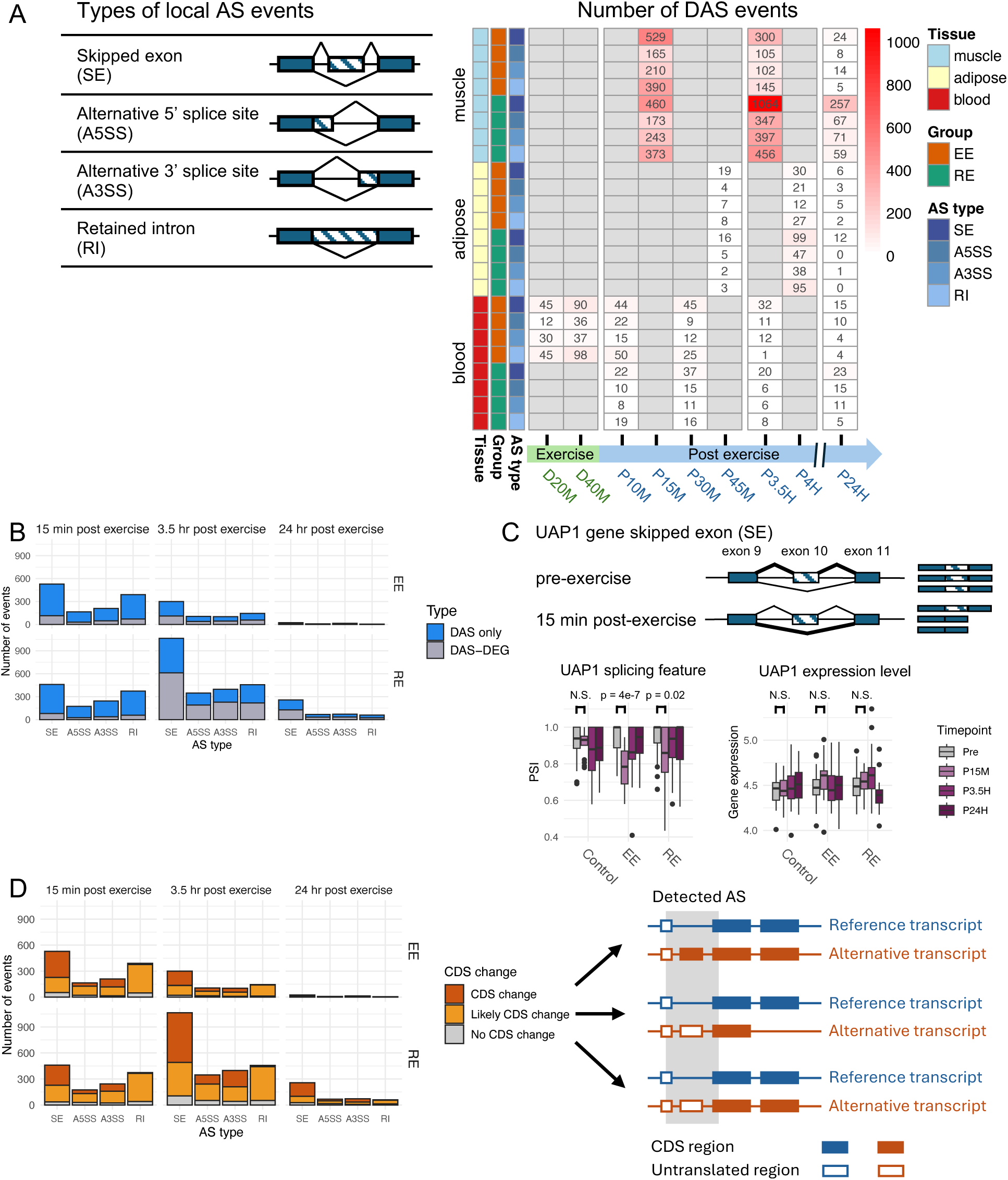
Differential AS events in SKM are mainly independent of differential gene expression and are linked to protein-coding changes. **A**, *Left*, Schematic of the major types of AS events analyzed in this study (SE, A5SS, A3SS, and RI). *Right*, Heatmap presenting the number of detected DAS events by group, tissue and AS event type. **B**, Bar graph showing the proportions of DAS events in SKM that were detected either in non-differentially expressed genes (DAS only) or in differentially expressed genes (DAS-DEGs), and their distribution by AS event type, time point, and exercise modality. **C**, Illustration of a representative DAS event in SKM. UAP1 exhibits a significant decrease in the PSI value of a SE event at exon 10 at P15M after EE, with no significant change in gene expression. T test, corrected p value < 0.05; N.S., not significant. **D**, Bar graph showing the proportions of SKM DAS genes associated with a protein-coding DNA sequence (CDS) change, a likely CDS change, or no CDS change. Provided to the right of the bar graph is a schematic describing each of the three types of changes associated with a detected AS event.

Comparison of the temporal patterns of DAS events across the three tissues revealed that the frequency of detected DAS events was considerably higher in SKM than in adipose tissue or blood, regardless of the exercise modality (**Fig. 2A, *Right panel***). In SKM, the number of DAS events in the early phase post-exercise (P15M) was comparable between the two exercise modalities (1294 for EE vs. 1249 for RE). However, while the number of DAS events decreased to 652 at the mid-phase post-EE (P3.5H), it increased to 2264 after RE. The same DAS events across time were more persistent after RE. A total of 84 DAS genes overlapped across time points after RE as compared to only 9 after EE (**Fig. S2**). Unlike SKM tissue, adipose tissue showed an increase in DAS events in both exercise modalities between the early and the mid-phase (from 38 to 90 for EE and 26 to 279 for RE), with overall more abundant DAS events following RE than EE. Conversely, blood exhibited a decrease in the number of DAS events between the early and the mid-phase in both exercise modalities (from 91 to 56 for EE and 79 to 40 for RE). In blood, the number of DAS peaked during the EE bout, with 132 and 261 events at D20M and D40M, respectively. DAS events dropped to the lowest levels at the late phase (P24H) across all three tissues. In SKM, DAS events declined from 652 at P3.5H to 51 at P24H for EE, and from 2264 to 454 for RE. The relative number of significant DAS events occurring at different timepoints in adipose and blood may have been influenced by differences in the number of samples obtained at some timepoints in these tissues. SKM sample numbers at all timepoints were comparable (see **Fig. 1B**). In all three tissues, SE and RI were the most prevalent DAS events.

### DAS genes and differentially expressed genes (DEGs) are largely non-overlapping

Because changes in transcription rate, also referred to as RNA Pol II elongation rate, represent one mechanism regulating AS (for review, see ^46,47^), we investigated the degree to which DAS events and differential gene expression occur at the same genes. We determined the number of genes undergoing DAS events only (DAS-only) and the number of genes undergoing both DAS events and differential gene expression (DAS-DEGs) at the different time points following each exercise modality. In adipose tissue and blood, the overall majority of DAS events at any timepoint or exercise modality occurred predominantly at genes showing no differential expression (**Fig. S3**). In SKM, the majority of DAS events detected in the early phase post-exercise for both modalities occurred at non-DEGs (∼82% DAS-only; **Fig. 2B**). The only exception was during the mid-phase post-RE, where SKM AS events were more common at DEGs (45% DAS-only after RE vs. 61% after EE). Taken together, these observations indicate that, with the exclusion of RE P3.5H in SKM, differential splicing and differential gene expression after exercise are largely independent.

As an example of acute exercise-regulated splicing in SKM, we show the analysis of UAP1, a gene encoding an enzyme involved in protein glycosylation that has been shown to promote antiviral responses ^48^. Among the AS isoforms derived from this gene, AGX2 corresponds to the canonical sequence, whereas AGX1 is generated by skipping protein-coding exon 10 ^49^. Previous enzymatic activity and structural analyses provide support that AGX1 and AGX2 have distinct catalytic properties due to differences in the active site architecture ^50,51^. We found that the PSI value of this AS event of UAP1 at chr1:162592735-162592782 was significantly decreased at P15M in the EE group, reflecting an increase in the presence of the spliced-out AGX1 isoform (**Fig. 2C**). However, the overall level of the UAP1 transcript was not altered by EE, indicating that regulation of this AS event was independent of the regulation of gene expression. Following RE, there was a borderline significant decrease in the PSI value for this AS event. Altogether, our data highlight distinctive patterns of regulation for UAP1 transcript isoforms and UAP1 gene expression with the regulated splicing consistently found in all P15M samples following EE.

### Most DAS events are associated with changes in the protein-coding sequence

To evaluate the potential influence of exercise-regulated AS on protein function in SKM, we examined the overall effects of DAS events on the protein-coding DNA sequence (CDS) of the corresponding genes. One limitation of using short-read RNA-seq and AS calling pipelines is that AS events involving the first or last exons are excluded from analysis because they cannot be quantified. CDS changes due to DAS were identified as either directly altering the annotated CDS (“CDS change” in **Fig. 2D**) or as being associated with an annotated transcript isoform that alters the CDS at a different AS site, usually involving the first or last exon (“Likely CDS change” in **Fig. 2D**, see Methods). A complete list of DAS events causing a CDS change or a likely CDS change are found in **Table S1**. Overall, 89% of the DAS events were associated with definite or likely CDS changes (**Fig. 2D** and **Table S1**). Furthermore, AS events that are not predicted to lead to changes in the CDS may also be functionally significant by altering RNA stability or post-transcriptional regulation. Similar observations were made in adipose tissue and blood (**Fig. S4**). These findings suggest that exercise-induced DAS events have a profound impact on protein composition.

### Enrichment of shared and distinct biological processes in DAS genes after EE and RE in SKM

Given the central role of SKM in mediating the effects of exercise and the large relative number of AS events detected, we compared the genes harboring DAS events and their pathway enrichment following EE and RE in this tissue. Paralleling the relative number of DAS events after EE and RE (see **Fig. 2A**), the number of genes affected by AS were comparable at P15M, but much greater in RE than EE at later timepoints (**Fig. 3A**). Half or more of the genes showing DAS following EE at P15M, P3.5H, and P24H also showed DAS with RE ((366, 263 and 17 genes, respectively). This suggested the existence of exercise-induced DAS gene-related programs that were either modality-specific or common to both modalities. The temporal trajectory of DAS events (**Fig. 2A**) and DAS genes (**Fig. 3A**) differed between EE and RE, with decreased changes observed at P3.5H after EE, and increased changes seen at P3.5H after RE.

**Figure 3.**
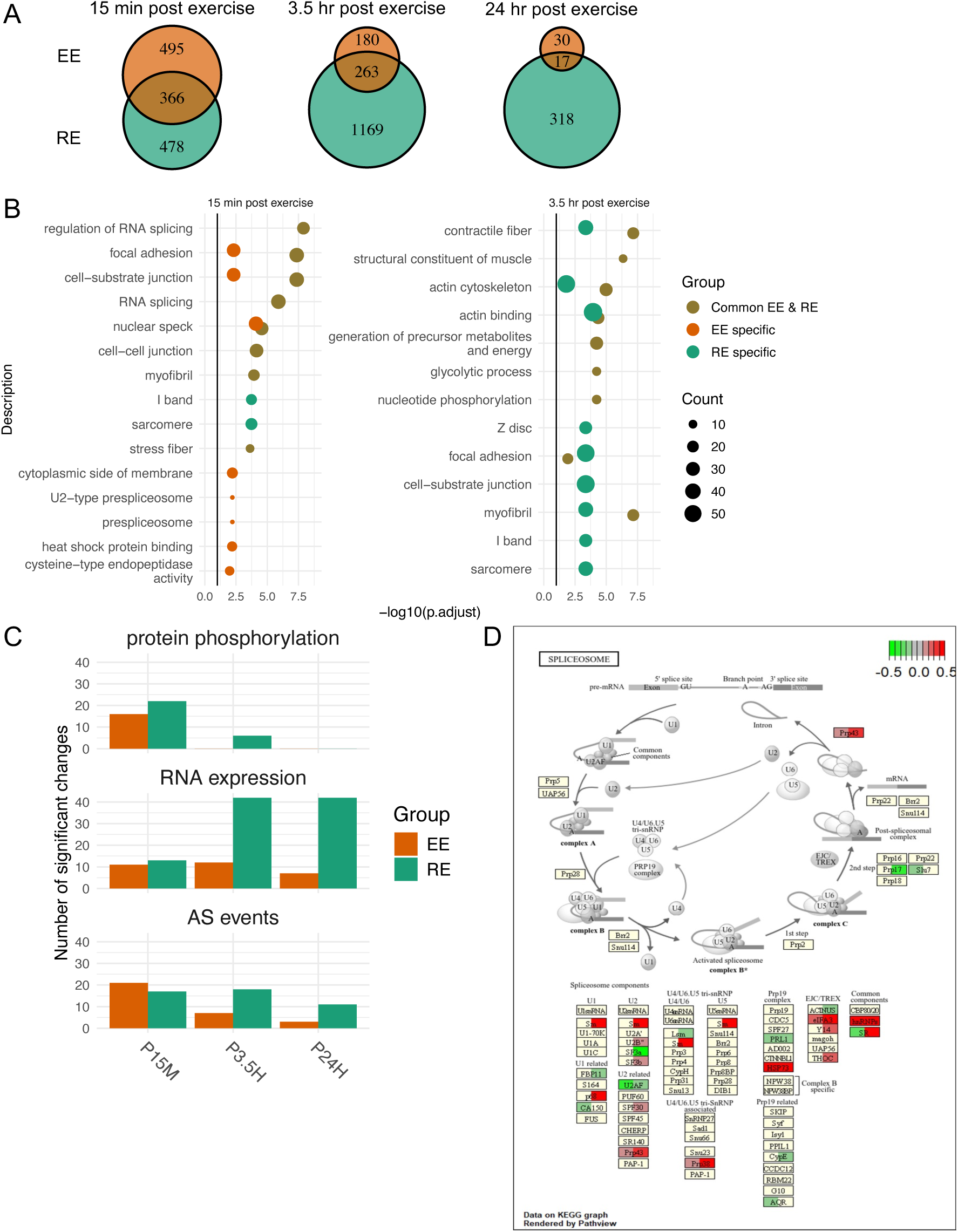
Exercise induces changes in spliceosome components at the protein phosphorylation, RNA, and AS levels in SKM. **A**, Venn diagram showing the overlap of SKM DAS genes between EE and RE at the three post-exercise time points. **B**, Pathway enrichment analysis on the SKM DAS genes identified at P15M and at P3.5H, with the DAS genes separated into 3 groups: DAS genes that are common to EE and RE, DAS genes that are EE-specific, and DAS genes that are RE-specific. The enrichment log10-adjusted p value is indicated on the X axis, and the dot size specifies the fold enrichment. **C**, Bar graph showing the number of significant changes observed among spliceosome-related genes at the level of protein phosphorylation, RNA expression, and AS at P15M, P3.5H, and P24H in each exercise modality. **D**, Spliceosome-related genes that were differentially expressed at P3.5H are highlighted in the KEGG Spliceosome Reference pathway (https://www.kegg.jp/pathway/map=map03040&keyword=spliceosome). Each spliceosome-related gene is symbolized by a box. Regulation of RNA levels by EE is represented on the half left of a gene box, while regulation of RNA levels by RE is on the half right. In the scale bar, red indicates the highest level of expression.

We performed GO enrichment analyses to determine which biological pathways were over-represented among i) DAS genes shared between EE and RE, ii) EE-specific DAS genes, iii) RE-specific DAS genes. At P15M, shared DAS genes were associated with regulation of RNA splicing, focal adhesion, and cell-substrate junction (**Fig. 3B**). Similarly to shared DAS genes, EE-specific DAS genes were enriched for nuclear speck, focal adhesion, and cell-substrate junction, and they additionally showed enrichment for heat shock protein binding. RE-specific DAS genes were enriched for sarcomere and I band, a region of the myofibril sarcomere containing actin, troponin, and tropomyosin. At P3.5H, shared DAS genes were enriched for contractile fiber, actin cytoskeleton, and myofibril, and RE-specific DAS genes were associated with actin binding, focal adhesion, and cell-substrate junction. No significantly enriched GO terms were found among EE-specific DAS genes at P3.5H (**Fig. 3B**). At P24H, RE-specific DAS genes were enriched for Z disc and muscle system process (**Fig. S5**). Although EE-specific and shared DAS genes at P24H showed significant enrichment for several processes, including acyl-CoA dehydrogenase activity and mitochondrial proton-transporting ATP synthase complex, respectively, only one or two DAS genes were annotated to these processes. Overall, shared as well as modality-specific DAS genes were enriched for processes related to RNA splicing and muscle structure or function at P15M and muscle structure or function at P3.5H.

### Spliceosome components are regulated by exercise at multiple levels in SKM

The pathway analysis described above indicated that in SKM, many DAS genes common to both exercise modalities were involved in the regulation of RNA splicing, suggesting that splicing machinery-related genes themselves may undergo AS in the early response to exercise. We assessed whether spliceosome components were also regulated by exercise at the protein phosphorylation and RNA expression levels. We focused on the 127 genes and proteins that have been identified as components of the pre-mRNA splicing process in the KEGG Pathway Database (https://www.genome.jp/kegg/pathway.html).

Using the phosphoproteomic data obtained by liquid chromatography tandem mass spectrometry (for details, see Methods and Pre-CAWG Landscape, **manuscript submitted**), we examined the phosphorylation status of 85 detectable spliceosome-related proteins, finding that 24 of them (28%) showed significant phosphorylation changes in samples from at least one timepoint after one exercise modality (**Fig. 3C, *Top***). Analysis of differential RNA expression and DAS among all 127 spliceosome-related genes resulted in the detection of 73 (57%) DEGs and 37 (29%) genes with DAS events in at least one timepoint and one exercise modality (**Fig. 3C, *Middle and Bottom***). The distribution of those DEGs across the splicing process by modality and timepoint is illustrated in **Fig. 3D and in Fig. S6**.

We also examined the occurrence of genes with highly correlated RNA expression profiles at baseline across all study participants using weighted gene co-expression network analysis (WGCNA; ^52^). Analysis of the pre-exercise SKM RNA-seq data identified a cluster/module of 300 genes that showed high enrichment for RNA splicing, RNA processing, RNA metabolic process, and RNA translation (**Fig. S7**). This WGCNA module was differentially regulated after exercise and included 42 spliceosome-related genes. These results suggest that a coherent module capturing inter-individual variation in the spliceosome transcripts is also regulated by exercise. On the whole, our data show that acute exercise modulates a large number of spliceosomal components at the protein phosphorylation, RNA expression, and RNA processing level. Out of the 127 spliceosome-related genes, a total of 84 (66%) demonstrate significant regulation with exercise, suggesting that a multi-layered modulation of the splicing process represents a major part of the response to acute exercise.

### Enriched splicing factor (SF) binding sites and SF regulation are associated with post-exercise DAS events

RBPs bind to specific motifs on pre-mRNA to regulate AS events (for review, see ^14^). Transcriptome-wide binding maps for human RBPs are compiled in the ENCODE database (https://www.encodeproject.org), having been identified using *in vivo* RBP crosslinking and immunoprecipitation (eCLIP) target profiling coupled with RNA-seq (eCLIP-seq; ^53,54^). To identify the RBPs having their target sites enriched at exercise-regulated AS loci, we evaluated the +/- 100 bp regions surrounding the splice sites of DAS events (see Methods). This analysis identified target site enrichment for a total of 49 RBPs at DAS sites for at least one time point and exercise modality at a Bonferroni corrected p value < 0.05 and a fold change > 1.5 (**Table S2**). After EE, the largest number of RBPs showed enrichment at P15M; fewer RBPs were enriched at P3.5H, and only one was enriched at P24H (**Fig. S8**). After RE, RBP enrichment was present at all timepoints. Presented in **Fig. 4A** are the RBPs showing the highest enrichment scores for each exercise modality and timepoint. Several RBPs were specific to an exercise modality. For example, the targets of RPS3, SF3A3 and UTP3 were enriched only at EE DAS regions, while the targets of EXOSC10 were enriched only at RE DAS regions.

**Figure 4.**
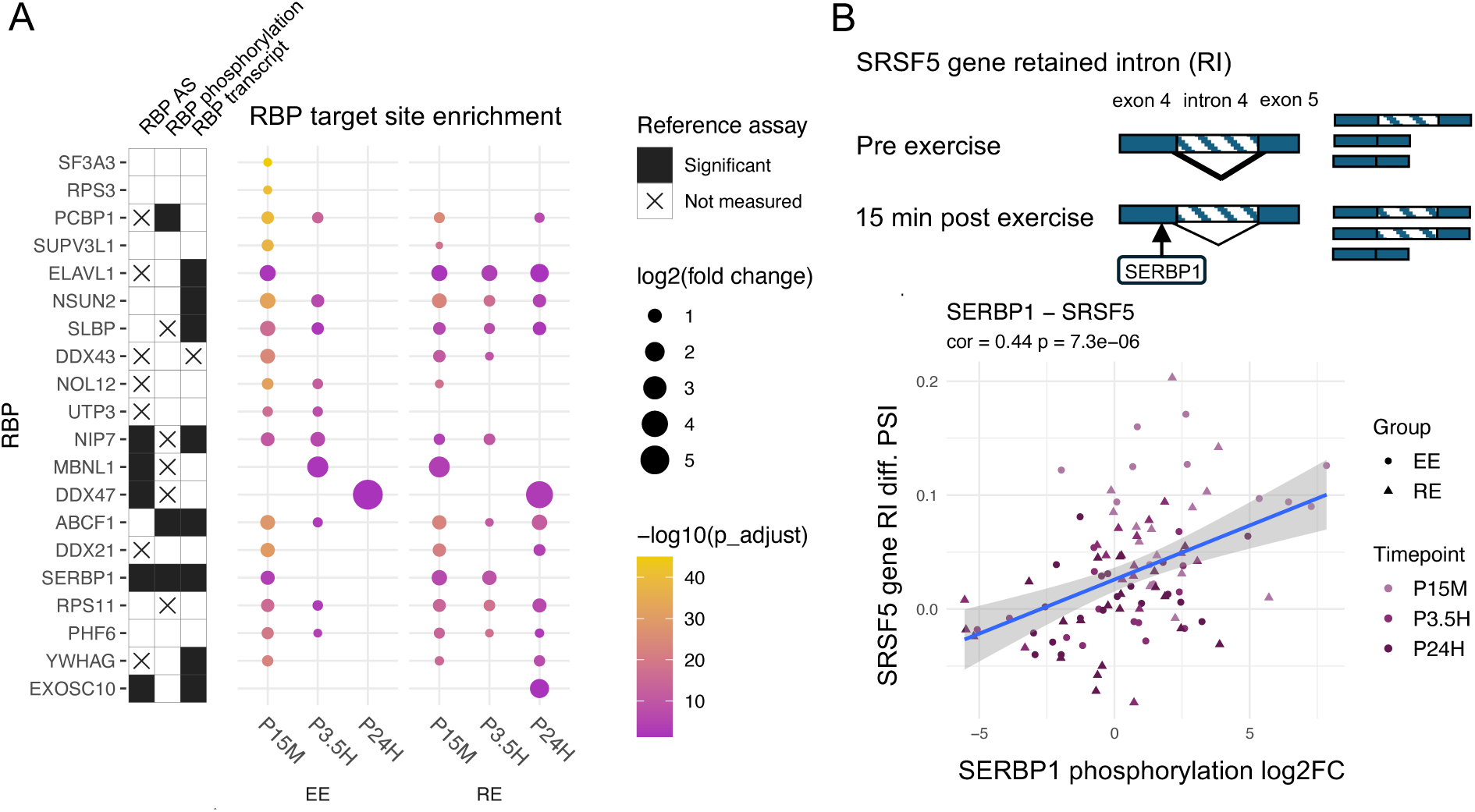
Implication of RBPs in the regulation of DAS events in SKM. **A**, RBP target sites enriched around DAS events. The time points for each exercise modality are indicated on the X axis. Provided on the Y axis are the RBP names, whether the RBP was differentially phosphorylated at one or more time points (solid bars), and whether the RBP-encoding gene was differentially expressed at one or more time points (solid bars). The color and size of circles represent the significance (log10-adjusted p value) of RBP target enrichment and the log2 fold change of RBP target enrichment, respectively. Top 4 RBPs by enrichment p values and top 4 RBPs by enrichment fold changes for each time point and each exercise modality were included in this plot. **B**, Example of a significant correlation between phosphorylation of a target-enriched RBP and the splice ratio (PSI value) of the target AS event. *Top*, Schematic of the DAS event occurring at P15M on the SRSF5 pre-mRNA and the binding of the RBP SERBP1. *Bottom*, Shown is the cross-sample correlation between the phosphorylation log2 fold change at SERBP1 Serine-330 compared to pre-exercise and the PSI value differences for the SRSF5 gene retained intron 4 compared to pre-exercise. Points are color-coded by time point and shaped by exercise modality, as indicated. The values of the correlation coefficient and the p value of the t test are indicated above the plot.

RBP activity can be modulated by its expression level, by phosphorylation, and by AS, which can alter its activity or its mRNA decay rate ^55–62^. We focused on the 20 RBPs showing the highest target enrichment at AS loci in SKM at each time point post-exercise. At the RNA expression level, 8 out of these 20 RBPs showed significant differential transcript expression (**Fig. 4A** and **Table S2**). While 16 of these 20 RBPs detected by mass spectrometry did not show significant changes in protein expression, few regulated proteins overall were detected after EE or RE (Pre-CAWG Landscape, **manuscript submitted**). We also found target-enriched RBPs (including 5 of the top 20 enriched RBPs) showing DAS at two or more time points following each exercise modality (**Fig. S9** and **Table S2**). Fifteen of the top 20 RBPs were detectable in phosphoprotein assays, and three showed differential phosphorylation at one or more post-exercise timepoints, namely SERBP1, ABCF1, and PCBP1. SERBP1 and ABCF1 transcripts were also upregulated (see **Table S2**). The differential phosphorylation sites of target-enriched RBPs, including SERBP1, ABCF1, and PCBP1, are presented in **Table S3**. We next examined the correlation between the phosphorylation of each RBP and the splice ratio (PSI values) of its individual target AS events across all participants from whom both the RBP phosphorylation and AS events were assayed (see Methods). A total of 823 RBP - AS event pairs involving 68 RBPs and 588 AS events showed significant correlations (**Table S4**). One example, shown in **Fig. 4B**, is the RBP SERBP1 and its AS target on the SRSF5 gene. The phosphorylation of SERBP1 at Ser-330 was highly correlated with the differential PSI value of a retained intron 4 AS event of SRSF5 (**Fig. 4B**). Overall, these findings lend support for the modulated activity of specific RBPs and their contribution to the programmatic regulation of AS following exercise.

### Implication of TF-driven circuits regulating post-exercise DAS events

Co-transcriptional modulation of splicing by DNA-binding TFs is an important contributor to the regulation of AS events ^19^. In the case of gene regulation, gene regulatory circuit triads (gene, TF, cis-binding site) can be inferred by detecting correlated TF activity, binding site accessibility changes, and gene expression changes ^63–65^. We applied a similar approach to identify putative TF-driven splice regulatory circuits (SRCs). An SRC contains differentially accessible chromatin sites that i) are distal or proximal to a DAS gene and/or in the DAS gene body, ii) are linked to a TF based on the mapping of its binding motifs, and iii) are correlated with a DAS event at the target gene (**Fig. 5A**). Leveraging the AS events and the chromatin accessibility peaks detected in the time-series after an acute exercise bout, we adapted the Multiome Accessibility Gene Integration Calling and Looping (MAGICAL) framework ^63^ to infer SRCs.

**Figure 5.**
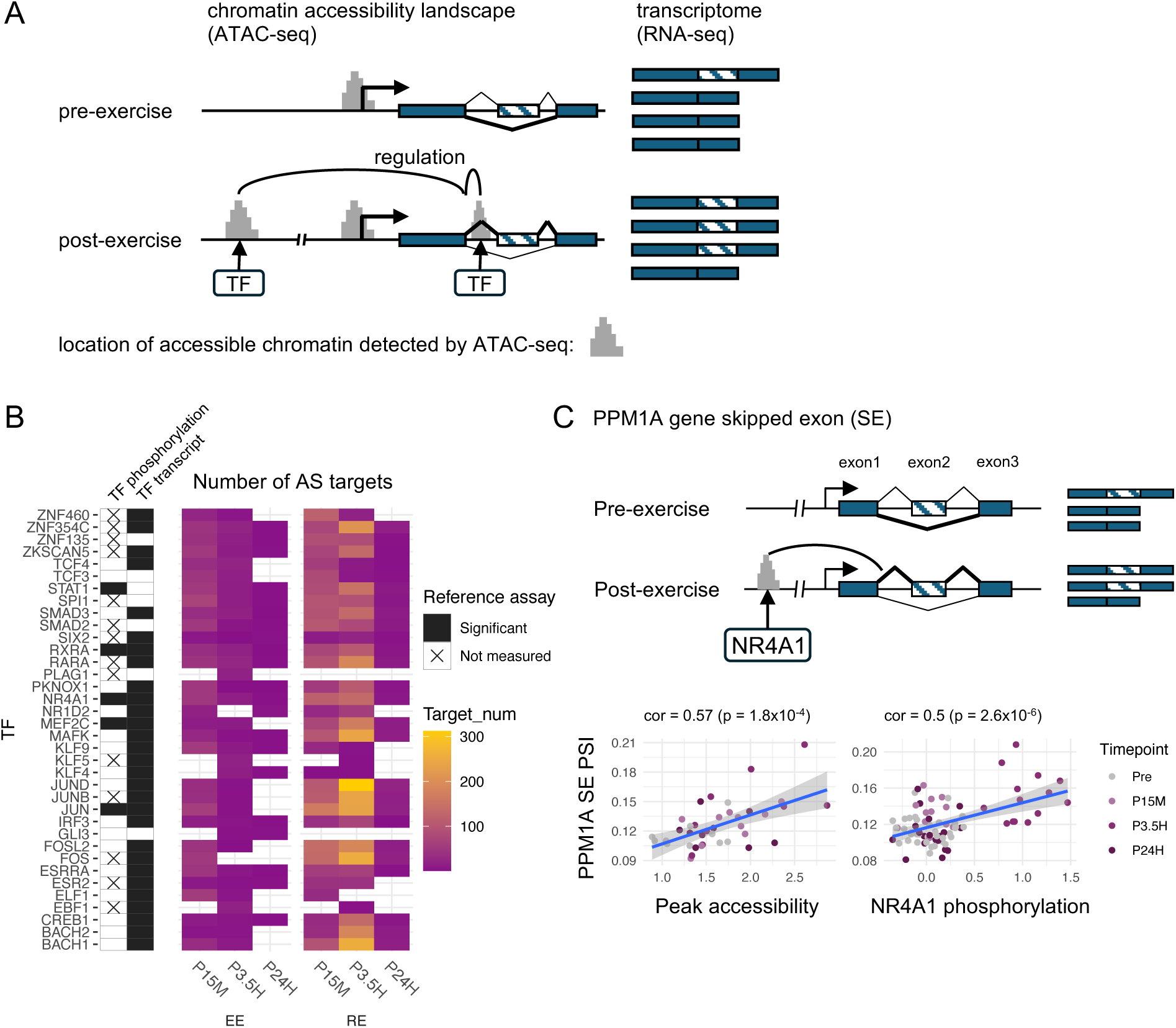
Implication of DNA-binding TFs in the regulation of DAS events in SKM. **A**, Schematic of the differential regulation of AS at the DNA level through altered chromatin accessibility and TF binding, identified using MAGICAL. Putative TFs are associated with differentially accessible chromatin regions that correlate with DAS events, forming splice regulatory circuits (SRCs). **B**, Shown are the top 10 TFs associated with the largest number of SRCs (i.e. the largest number of target DAS events) identified for each time point and each exercise modality using MAGICAL. The time points for each exercise modality are indicated on the X axis. Provided on the Y axis are the names of the TFs; also denoted on the Y axis is whether the TF was differentially phosphorylated at one or more time points (solid bars), and whether the TF-encoding gene was differentially expressed at one or more time points (solid bars). The number of target AS events is color-coded, as indicated. **C**, Example of a significant SRC involving the binding of NR4A1 to the PPM1A gene and correlated with a SE event at exon 2. *Top*, Schematic of the SE event occurring on the PPM1A pre-mRNA and the binding of the TF NR4A1 to a region located ∼75 kb upstream from the splice site of PPM1A gene. *Bottom*, Shown is the correlation between peak accessibility at the upstream region of the PPM1A gene and the PSI values for the SE event across time points in RE (*Left*), and the correlation between NR4A1 phosphorylation at Serine-14 and the PSI values for the SE event across time points in RE (*Right*). Points are color-coded by time point, as indicated. The values of the correlation coefficient and the p value of the t test are indicated above the plot.

MAGICAL detected multiple differentially accessible chromatin regions containing motifs for putative TFs in SKM, which correlated with DAS events following either EE or RE. Overall, we inferred a total of 2764 SRCs, 46% of which were associated with regulated AS in the absence of regulation of gene expression (**Table S5**). The TFs associated with the largest number of SRCs are shown in **Fig. 5B**. Top candidate TFs in both exercise modes included the immediate early genes FOS, JUN, JUNB, FOSL2, NR4A1, as well as STAT1, MEF2C, RXRA, RARA, and SMAD3.

TF activity can be regulated by its expression level and by post-translational modifications such as phosphorylation ^66,67^. We assessed the RNA expression and protein phosphorylation levels of the main TFs implicated in regulated SRCs at each time point after acute exercise. The majority (29 out of 36) showed differential transcript levels at one or more time points in each exercise modality, and a few (JUN, MEF2C, NR4A1, STAT1, and RXRA) exhibited significant changes at both the RNA and phosphorylation levels. Notably, we determined that JUN was differentially phosphorylated in the early phase post-RE, whereas NR4A1 was differentially phosphorylated in the mid-phase (**Table S6**). We illustrate a TF-driven SRC inferred through integrative analysis of AS and chromatin accessibility patterns. In the AS-regulated *PPM1A* gene, which is not regulated at the level of gene expression, the accessibility of a chromatin region located ∼75 kb upstream of a SE at exon 2 was strongly correlated with the inclusion of this exon, suggesting a specific co-transcriptional AS regulation mechanism. NR4A1, which was identified as the putative TF binding to that open chromatin region, also showed coordinated phosphorylation patterns (**Fig. 5C**). These correlated changes in site accessibility, TF phosphorylation, and DAS in the absence of gene expression changes provides strong support for NR4A1 binding mediating the effects of exercise on AS of the *PPMIA* gene.

## Discussion

Accumulating evidence suggests that AS is a crucial mechanism for promoting transcriptome and proteome diversity and that AS is modulated by diverse mechanisms. We uncovered the dynamics of the AS landscape following two modalities of acute exercise in SKM, blood and adipose tissue. Through integrated analyses of transcriptomic, epigenomic, and phosphoproteomic data derived from the same subjects, we inferred regulatory mechanisms underlying those AS changes in response to exercise. We found that acute exercise induced substantial changes in mRNA splicing, with the majority of these events altering protein composition. The patterns of AS alterations showed considerable differences following EE and RE. The regulation of AS occurred largely independently of regulation of gene expression. In SKM, which features the largest number of detected DAS events, the exercise-regulated AS landscape was associated with RNA splicing and muscle biology-related processes. The involvement of specific RBPs and DNA-binding TFs in directing AS events was inferred from the multi-omic data, providing insight into the underlying mechanisms mediating exercise-regulated AS.

We found that components of the spliceosome complex, which plays a central role in AS by removing intronic regions from mRNA precursors and joining exons to generate mature mRNAs, were highly regulated by exercise at the levels of RNA expression, RNA splicing, and protein phosphorylation. Furthermore, the enrichment of shared and modality-specific DAS genes for splicing regulation-related terms such as nuclear speck are consistent with modulation of the subcellular localization of SFs, as previously described ^68^. These findings are consonant with the formulation that external exercise acts via signal transduction pathways to affect the splicing machinery, thereby resulting in altered AS patterns, and that alterations in the splicing machinery may include changes in the phosphorylation/activity, expression, and localization of SFs, as well as interactions with the transcription machinery ^68^. Consistent with our findings, a recent mouse study demonstrated that endurance exercise regulates mRNA splicing and spliceosome-related nuclear proteins in SKM ^69^. We also applied WGCNA to the SKM pre-exercise transcriptome data. WGCNA identifies gene modules based on transcript group co-regulation across individuals. Among the modules that we identified through WGCNA analysis of the pre-exercise transcriptome data, MEpurple was highly enriched in spliceosomal genes. Many of those spliceosomal genes, which were identified only from pre-exercise individual variation in expression, were also regulated after either EE or RE (see **Fig. S7** and **Fig. 3C**). Thus, this WGCNA module may capture a transcriptome spliceosome set point that varies among individuals and that is coherently regulated by exercise. This analysis raises the possibility that the spliceosome set point represents another factor that may contribute to inter-individual variations in exercise capabilities and responses ^11^. Additionally, we found that DAS events affected many spliceosome genes (see **Fig. 3C**), suggesting that exercise regulation of splicing extends to remodeling the spliceosome complex itself.

As summarized in **Fig. 6**, we employed an integrative analysis of the multi-omics data to further investigate the molecular mechanisms underpinning the AS changes induced by exercise. The two exercise modalities studied, EE and RE, showed differences in the genes and splice junctions associated with DAS events, as well as in the overall temporal pattern of regulated splicing. EE and RE induced both common and modality-specific DAS genes in SKM (see **Fig. 3A**). EE showed slightly more DAS genes at P15M, whereas RE showed more than thrice as many DAS genes at P3.5H and nearly 9 times more DAS genes at P24H than EE. Additionally, the pathways associated with EE-vs. RE-specific DAS genes differed (see **Fig. 3B**). Notably, the over-representation of heat shock protein binding at P15M solely among EE-specific DAS genes may reflect modality-divergent effects on heat shock responses, as suggested by the MoTrPAC Study Group (Pre-CAWG Landscape, **manuscript submitted**) and the previously described modulatory effect of acute exercise on heat shock protein expression ^70,71^. Consonant with previous reports that gene expression and splicing mediate distinct biological processes ^72^, overall, a majority of exercise-induced DAS events occurred at genes that showed no differential expression. In contrast, however, after RE at P3.5H, the majority of DAS genes were also DEGs (see **Fig. 2B**). Notably, in SKM, the number of DEGs described by the MoTrPAC Study Group was highest at P3.5H after RE relative to EE or any other timepoint (Pre-CAWG Landscape, **manuscript submitted**). Altered transcript elongation rate is one contributor to the regulation of AS (for review, see ^47^). Thus, one possible explanation for a higher overlap of DAS and DEG at RE P3.5H could be changes in the relative contribution of increased elongation rate and other splicing mechanisms to AS at RE P3.5H.

**Figure 6.**
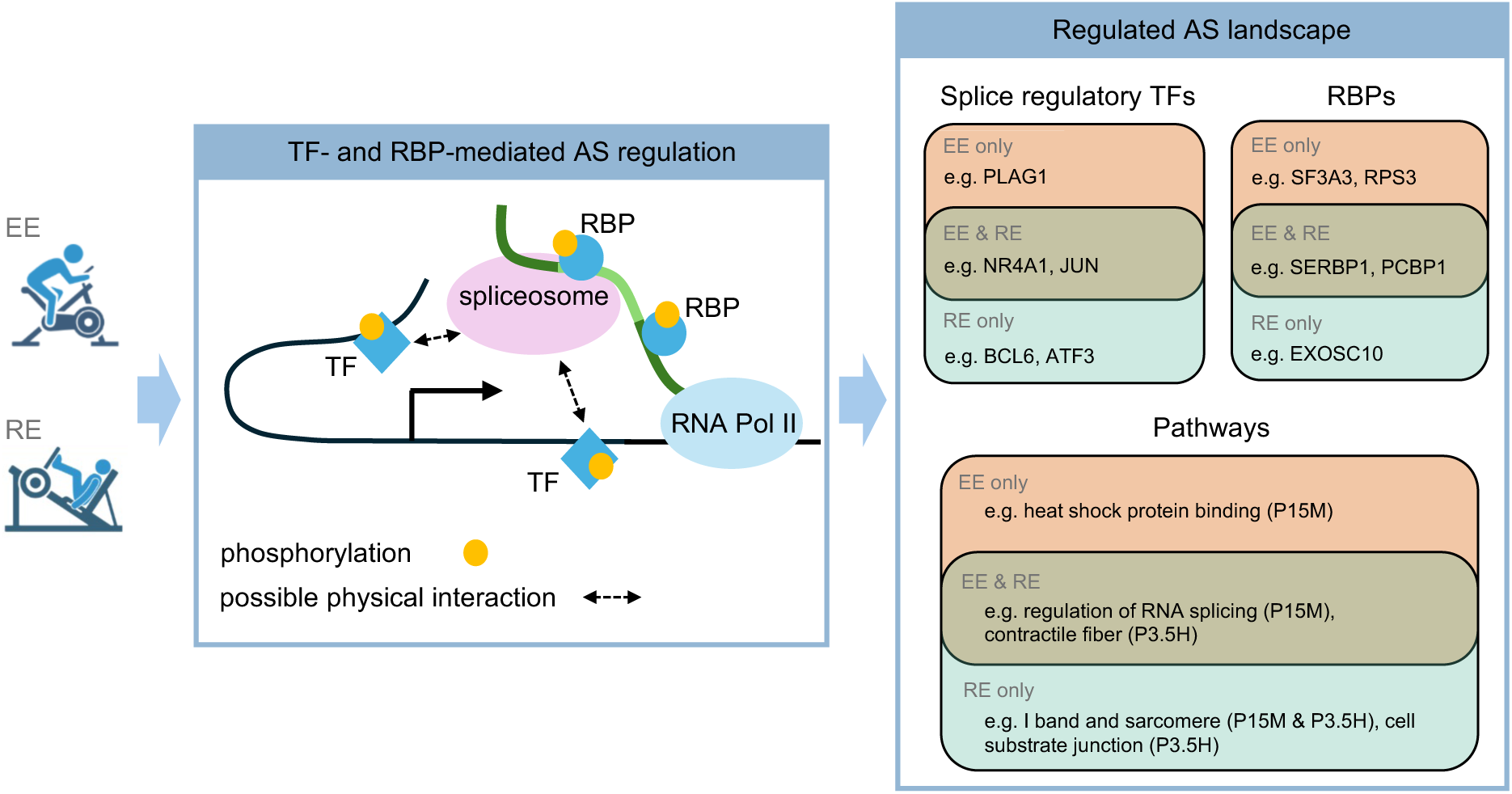
Model of acute exercise-induced modulation of AS in SKM and its regulatory mechanisms. *Left*, The AS landscape is modulated by EE and RE via DNA-binding TF-dependent and RBP-dependent mechanisms, along with changes in the phosphorylation state of those TFs and RBPs. *Right*, Illustration of the exercise-regulated landscape through examples of modality-specific and shared TFs and RBPs, along with the biological pathways enriched among modality-specific and shared DAS genes.

Enrichment analysis of the pattern of RBP targets at DAS event junctions implicated candidate RBPs in splicing regulation (see **Fig. 4A**). Altered RBP gene expression and/or phosphorylation was found for many of these RBPs. As we did not detect concomitant changes in protein expression for any RBP, we were unable to definitely determine whether the changes in RBP gene expression influenced the immediate splicing responses seen after exercise. We note, however, that overall few proteomic changes were detected following exercise. In SKM, for example, a total of 7917 DEGs were detected after RE, whereas only 31 proteins were significantly regulated; with EE, there were altogether 4042 DEGs and only 4 differentially regulated proteins ([see Fig. 2A in] Pre-CAWG Landscape, **manuscript submitted**). Thus, as generally observed in proteomics due to a variety of factors, the mass spectrometry assays may have been relatively insensitive to changes in RBP protein expression ^73–75^. Alternatively, any effects of regulation of transcription may be delayed beyond the study period. We also detected DAS events for several of the enriched RBPs, which may regulate RBP level or activity. Overall, the overlap of RBP phosphorylation changes, DEGs, and DAS events at RBP-encoding genes for RBPs showing target enrichment at DAS regions provides further support for the role of specific RBPs in mediating the regulation of AS by exercise.

There is increasing interest in the role of RBP modulation in mediating exercise responses ^38,76^. Alterations in RBP expression or activity have been implicated in many biological processes including inflammation, cell survival and cell migration. Among the candidate RBPs that showed differential phosphorylation in this study, PCBP1 is considered a typical RBP that harbors three RNA-binding hnRNP K homology (KH) domains ^77^, regulates splicing, and has been previously implicated in SKM differentiation and development ^78^. In contrast with PCBP1, the target-enriched RBPs SERBP1 and ABCF1 lack well-characterized RNA-binding domains ^53^. SERBP1 contains RG/RGG repeats. RBPs with these repeat regions are essential for brain function and have been involved in neurological and neuromuscular diseases ^79^. SERBP1 is known to enhance the stability of SERPINE1/PAI1 mRNA ^80^. ABCF1 has been involved in the regulation of mRNA translation ^81^. The differentially regulated phosphosites that we identified at one or more time points in ABCF1 (Ser-109, Ser-140, Thr-108) and PCBP1 (Ser-246) were previously linked to changes in protein activity, consistent with the modulation of their activity by acute exercise. In ABCF1, phosphorylation at Ser-109 and Ser-140 by CK2 has been implicated in the association of the phosphoprotein binding partner eIF2 with 80S ribosomal and polysomal fractions ^82^, while Thr-108 may affect the function of the adjacent CK2-phosphorylated Ser-109 ^83^. Functional studies of PCBP1 previously showed that Ser-246 phosphorylation is involved in the upregulation of a target gene (MOR; ^84^). The exercise-regulated Ser-330 phosphosite that we detected in SERBP1 has been previously reported to be exercise-regulated ^85^. The effects of this phosphosite on SERBP1 activity is unknown. ABCF1 has not been linked to muscle physiology. Further investigation will be needed to corroborate the functional roles of the RBPs we identified, including PCBP1, SERBP1 and ABCF1, in mediating the modulatory effects of exercise on AS.

Besides altering gene expression, DNA-binding TFs can regulate pre-mRNA splicing and consequently transcript isoform expression. Our integrated analysis of chromatin accessibility, RNA expression, and AS enabled us to infer SRCs in the context of acute exercise. In the present work, we uncovered exercise-associated SRCs formed by DAS genes and differentially accessible chromatin regions containing motifs for putative TFs. Furthermore, the majority of those putative TFs exhibited differential gene expression, suggesting that they are regulated by acute exercise. In keeping with this, JUNB, FOSL2, NR4A1, and SMAD3 were formerly reported to be induced by acute exercise in SKM ^9,86^. A portion of the candidate TFs were also differentially phosphorylated at one or more time points following EE or RE.

TFs associated with the largest number of SRCs, which concomitantly showed alterations in their RNA level and in their phosphorylation state, comprised JUN, MEF2C, NR4A1, and RXRA. Consistent with the functional roles of MEF2C and STAT1 during exercise, an increase in the DNA-binding activity of MEF2C in SKM was previously described following acute exercise ^87^, and activation of the IL-6/STAT1/STAT3 signaling pathway was reported in rat SKM following acute RE, including an increase in STAT1 phosphorylation ^88^. We observed altered MEF2C phosphorylation at Ser222 at P15M after RE. A cancer-related study demonstrated that Ser222 phosphorylation is necessary for activating MEF2C-mediated gene expression program, implying that it affects MEF2C transcriptional activity ^89^. STAT1, which exhibited differential phosphorylation at Ser727 at P15M after RE, was also shown to depend on Ser727 phosphorylation for optimal transcriptional activity ^90^. Phosphorylation of JUN at Ser243 was previously reported to negatively regulate its DNA-binding activity (see ^91–93^), supporting the role of JUN activity in the regulation of AS during exercise. Similarly, phosphorylation of RXRA at Ser260 is thought to inhibit its function through decreased DNA binding and/or possibly reduced transcriptional activity ^94–96^. Moreover, differential phosphorylation of RXRA at Ser21 was identified at both P15M and P3.5H after RE, and murine studies suggest that Ser21 phosphorylation may affect RXRA function only in the presence of its ligand, retinoic acid ^97,98^. While there is no direct evidence of the role of Ser14 phosphorylation in NR4A1, earlier reports indicate that this residue is in the N-terminal transactivation domain and that its phosphorylation might contribute to transactivation activity ^99,100^ and possibly nuclear localization of NR4A1 ^101^. Taken together, these previous reports support the view that acute exercise may affect MEF2C, RXRA, STAT1, and NR4A1 activity via phosphorylation changes.

### Study limitations

The short-read RNA-seq-based methods used in our analysis did not allow detection of alternative transcription start sites (ATSS) and alternative transcription termination sites (ATTS; also referred to as alternative polyadenylation sites or APA), even though they are known to contribute significantly, alongside AS, to transcript isoform diversity ^102^. Our DAS analysis relied on computational methods that may have some advantages and drawbacks compared to other analysis tools. We used a combination of DARTS/rMATS-turbo and SUPPA2, both of which are event-based methods, in contrast with isoform-based approaches such as Cufflinks and DiffSplice, and exon-based approaches like DEXSeq and limma (see ^103^). Based on a previous systematic evaluation, rMATs along with exon-based methods performed generally well, while SUPPA2 detected few DAS genes and showed a high false discovery rate. Future studies using long-read RNA-seq methods will be useful in expanding detection of regulated mRNA isoforms by exercise. Genetic loss-of-function experiments will be needed to validate the role of specific SFs in mediating the AS response to acute exercise. While previous work reported the impact of aging and nutrition on AS in human SKM ^104^, we were unable to examine the effects of aging or nutrition on AS, given the limited size of our study cohort and the overnight fast of all participants prior to the intervention.

### Conclusions

We present a characterization of the AS landscape of the human response to acute EE and RE in multiple tissues using a multi-omic, integrative analysis approach. Our investigation suggests that the AS alterations induced by acute exercise play a functional role in the tissue responses to exercise. We identify the putative regulatory mechanisms mediating exercise-regulated AS. Besides contributing to a better understanding of the dynamic molecular and biological changes occurring with exercise, this work could help to develop personalized exercise therapy in the treatment of chronic disorders such as metabolic, neurological, cardiovascular diseases, and cancer. Overall, we provide insight into the global regulation of AS by exercise and its underlying mechanisms, and our study supports a significant role for AS in mediating the tissue responses to acute exercise.

## Supplemental information

### Figures S1-S9

**Figure S1.** Schematic illustrating the calculation of PSI values from RNA-seq reads. *Top*, rMATS uses the number of junction reads to quantify the transcripts including the alternative regions and the transcripts excluding the alternative regions, and calculate the PSI values based on these quantifications. *Bottom*, SUPPA uses all reads and quantifies distinct transcripts by pseudo-alignment of reads to annotations, and calculated PSI values based on the TPM values of quantified transcripts.

**Figure S2.** Overlap between SKM DAS genes across time points within each exercise modality.

**Figure S3. A**, Bar graph showing the proportions of adipose tissue genes associated with DAS events (“DAS genes”) that correspond to either non differentially expressed genes (DAS only) or differentially expressed genes (DAS-DEGs), and their distribution by AS event type, time point, and exercise modality. **B**, Same as **A** for blood genes. Related to Fig. 2B.

**Figure S4.** Bar graph showing the proportions of adipose tissue DAS genes associated with a protein-coding sequence (“CDS”) change, a likely CDS change, or no CDS change. Provided to the right of the bar graph is a schematic describing each of the three types of changes resulting from a detected AS event. Related to Fig. 2D.

**Figure S5.** Pathway enrichment analysis on the SKM DAS genes identified at P24H, with the DAS genes separated into 3 groups: DAS genes that are common to EE and RE, DAS genes that are EE-specific, and DAS genes that are RE-specific. Related to Fig. 3B.

**Figure S6.** Spliceosome-related genes that were differentially expressed at P15M and at P24H are highlighted in the KEGG Spliceosome Reference pathway (https://www.kegg.jp/pathway/map=map03040&keyword=spliceosome). Each spliceosome-related gene is symbolized by a box. Regulation by EE is represented on the half left of a gene box, while regulation by RE is on the half right. In the scale bar, red indicates the highest level of expression. Related to Fig. 3C.

**Figure S7.** Weighted gene co-expression network analysis (WGCNA) of the RNA-seq data from pre-exercise SKM samples identifies a spliceosome-related module.

**Figure S8.** Bar graph showing the number of RBPs enriched at DAS sites per time point for each exercise modality in SKM.

**Figure S9**. Bar graph showing the number of RBPs enriched at DAS sites per time point for each exercise modality in SKM, which concurrently underwent DAS.

### Tables S1-S6

**Table S1. List of the DAS events and their association with CDS changes, likely CDS changes and no CDS changes.**

**Table S2. List of the putative RBPs implicated in DAS events.** The 54 RBP candidates were identified by RBP target enrichment analysis over the +/- 100 bp region surrounding the splice site of a detected DAS event (see Methods). A significant enrichment is above Bonferroni corrected p value < 0.05 and FC > 1.5.

**Table S3. List of target-enriched RBPs that were differentially phosphorylated following either EE or RE.**

**Table S4. List of RBP - AS event pairs.** A pair is defined based on significant correlation between the phosphorylation of an RBP and the splice ratio (PSI value) of its individual target AS event across all samples that were assayed for RBP phosphorylation and AS events (see Methods).

**Table S5. List of splice regulatory circuits (SRCs) that were predicted using MAGICAL.** Specified is whether an SRC is associated with gene expression regulation.

**Table S6. List of the main TFs driving SRCs that were differentially phosphorylated following either EE or RE.**

## Resource availability

### Lead contact

Further information and requests for resources and reagents should be directed to and will be fulfilled by the lead contact, Stuart Sealfon.

### Materials availability

This study did not generate new unique reagents.

### Data and code availability

The *MotrpacHumanPreSuspensionAnalysis* R package (https://github.com/MoTrPAC/MotrpacHumanPreSuspensionAnalysis) contains functions to generate visualizations, as well as data objects for differential abundance analysis, summary statistics for normalized expression data, feature to gene mapping, and molecular signature datasets as described in the methods.

The MotrpacHumanPreSuspensionData R package (https://github.com/MoTrPAC/MotrpacHumanPreSuspensionData) is a package containing individual level data, which cannot be made openly available and can be accessed with approval from a data access committee. See motrpac-data.org for details.

Data used in the preparation of this article were obtained from the Molecular Transducers of Physical Activity Consortium (MoTrPAC) database, which is available for public access at motrpac-data.org. The specific version released is version 1.3 of the human precovid sed adu.

Lastly, the precovid-analyses GitHub repository (https://github.com/MoTrPAC/precovid-analyses) contains code and individual parameters for each figure panel, utilizing the MotrpacHumanPreSuspensionAnalysis and MotrpacHumanPreSuspensionData packages.

## STAR Methods

### Experimental model and study participant details

#### IRB

The Molecular Transducers of Physical Activity Consortium (MoTrPAC) (NCT03960827) is a multicenter study designed to isolate the effects of structured exercise training on the molecular mechanisms underlying the health benefits of exercise and physical activity. Methods are described here in sufficient detail to interpret results, with references to supplemental material and prior publications for detailed descriptions. The present work includes data from prior to the suspension of the study in March 2020 due to the Covid-19 pandemic. The study was conducted in accordance with the Declaration of Helsinki and approved by the Institutional Review Board of Johns Hopkins University School of Medicine (IRB protocol # JHMIRB5; approval date: 05/06/2019). All MoTrPAC participants provided written informed consent for the MoTrPAC study indicating whether they agreed to open data sharing and at what level of sharing they wanted. Participants could choose to openly share all de-identified data, with the knowledge that they could be reidentified, and they could also choose to openly share limited de-identified individual level data, which is lower risk of re-identification. All analyses and resulting data and results are shared in compliance with the NIH Genomic Data Sharing (GDS) policy and DSMB requirements for the randomized study.

#### Participant Characteristics

Volunteers were screened to: (a) ensure they met all eligibility criteria; (b) acquire phenotypic assessments of the study population; and (c) verify they were healthy and medically cleared to be formally randomized into the study (cite Clinical Landscape) ^105^. Participants were then randomized into intervention groups stratified by clinical site and completed a pre-intervention baseline acute test. A total of 176 participants completed the baseline acute test (EE=66, RE=73, and CON=37), and among those, 175 (99%) had at least one biospecimen sample collected. Participants then began 12 weeks of exercise training or control conditions. Upon completion of the respective interventions, participants repeated phenotypic testing and the acute test including biospecimen sampling. Due to the COVID-19 suspension, only 45 participants (26%) completed the post-intervention follow-up acute exercise test (some with less than 12 weeks of training), with 44 completing the post-intervention biospecimen collections. See Clinical Landscape for further detail.

#### Participant Assessments

As described in Clinical Landscape and protocol paper, prior to randomization, participant screening was completed via questionnaires, and measurements of anthropometrics, resting heart rate and blood pressure, fasted blood panel, cardiorespiratory fitness, muscular strength, and free-living activity and sedentary behavior (cite Clinical Landscape) ^105^. Cardiorespiratory fitness was assessed using a cardiopulmonary exercise test (CPET) on a cycle ergometer (Lode Excalibur Sport Lode BV, Groningen, The Netherlands) with indirect calorimetry. Quadriceps strength was determined by isometric knee extension of the right leg at a 60° knee angle using a suitable strength device ^105^. Grip strength of the dominant hand was obtained using the Jamar Handheld Hydraulic Dynamometer (JLW Instruments, Chicago, IL). See Clinical Landscape for participant assessment results.

#### Acute Exercise Intervention

The pre-intervention baseline EE acute bouts (**Fig. 1A**) were composed of three parts: (1) a 5min warm up at 50% of the estimated power output to elicit 65% VO2peak, (2) 40min cycling at ∼65% VO2peak, and (3) a 1min recovery at ∼25 W. The pre-intervention baseline RE acute bouts were composed of a 5min warm-up at 50-60% heart rate reserve (HRR) followed by completion of three sets each of five upper (chest press, overhead press, seated row, triceps extension, biceps curl) and three lower body (leg press, leg curl, leg extension) exercises to volitional fatigue (∼10RM) with 90sec rest between each set. Participants randomized to CON did not complete exercise during their acute tests. Participants rested supine for 40min to mirror the EE and RE acute test schedule. See Clinical Landscape for acute bout results.

#### Biospecimen Collection

To standardize conditions prior to the acute test, participants were instructed to comply with a variety of controls related to COX-inhibitors, biotin, caffeine, alcohol, exercise, and final nutrition consumption ^105^. Blood, muscle, and adipose samples were collected for the acute test at specific timepoints before, during, and after exercise (**Fig. 1A**). See Multi-omic Landscape for the details of multi-omic coverage for each tissue.

Participants arrived fasted in the morning, and rested supine for at least 30min prior to pre-exercise biospecimen collections. All participants had pre-exercise sampling from all three tissues. After the pre-exercise biospecimen collections, the subsequent collection timepoints varied for each training group and their randomized temporal profile (early, middle, late, all). Temporal profiles were used to reduce the burden of repeat sampling while still maintaining biochemical coverage of the full post-exercise period. During and after exercise, biospecimen collection varied by randomized group, tissue, and temporal group. At 20 and 40 minutes of exercise, blood was collected from only EE and CON groups due to limitations in safely obtaining blood during RE. Immediately post exercise, at 10 minutes, a blood sample was collected in all three groups. Finally, samples from all three tissues were collected at early (SKM at 15, blood at 30, and AT at 45 minutes post-exercise), intermediate (blood and SKM at 3.5, and AT at 4 hours post-exercise), and late (24 hours post-exercise in all tissues) timepoints. The post-exercise timepoint began at the completion of the 40 minutes of cycling, third set of leg extension, or 40 minutes of rest, depending on the randomized group. Except for the EE blood collection timepoints during exercise, participants rested supine or seated for biospecimen collections. The collection and processing protocols for each tissue were previously described in the human clinical protocol design paper ^105^. See Clinical Landscape for biospecimen collection success results and other details.

### Molecular Quantification Methods

#### ATAC-sequencing

ATAC-seq (Assay for Transposase Accessible Chromatin using sequencing) assays were divided by tissue, with SKM tissue assayed at Icahn School of Medicine at Mount Sinai and blood PBMCs assayed at Stanford University. Samples for ATAC PBMC processing were randomized as described in https://github.com/MoTrPAC/clinical-sample-batching. SKM samples were processed for ATAC-seq by study group (CON, EE, and RE). Eight to 12 samples were processed at a time for nuclear extraction and library preparation, and all the samples in the study group were sequenced together. For muscle, nuclei from aliquoted tissue samples (10 mg for Muscle tissues) were extracted using the Omni-ATAC protocol with modifications. Two tissue-specific consortium reference standards were included for sample processing QC. 500-600 ul 1X homogenization buffer with protease Inhibitor was added to the sample micronic tubes plus three 1.8 mm ceramic bead media. Tubes were vortexed for 5-10 seconds 3 times and kept on ice. The mixture was homogenized using Bead Ruptor Elite (Bead Mill Homogenizer) and Omni BR Cryo Cooling Unit at the settings: Tube 1.2 ml, Speed: 1.0, Cycles: 2, Time: 20 Sec, Dwell Time: 20 Sec, Temperature: 4-8⁰C. The homogenized mixture was filtered with Mini 20 um Pluristrainer and waited for 20-30 seconds. The homogenate tubes were centrifuged with filters attached briefly if the homogenate did not pass through the filter easily. Nuclei were stained with Trypan blue dye 0.4% and counted using Countess II Automated Cell Counter. Depending upon the counts, the number of nuclei to use for transposition was calculated. For Human Pre-covid muscle samples, 250K-300K nuclei were used. Calculated nuclei were added to 1 ml RSB in 2.0 ml Eppendorf tubes, mixed well and centrifuged at 510 RCF for 10 min on fixed angle centrifuge at 4⁰C. 70% supernatant was removed without disturbing pellet at the bottom and nuclei were again centrifuged at 510 RCF for 5 minutes. Almost all of the supernatant was removed. A 50 μl transposition mix was added to the pellet, and mixed by pipetting 6 times, then incubated at 37⁰C on a thermomixer with 1000 rpm for 30 minutes and immediately transferred to ice after incubation. The transposed DNA was purified using Zymo DNA Clean and Concentrator kit (Zymo research, D4014). The DNA product was amplified using NEBnext High-Fidelity 2x PCR Master Mix (NEB, M0541L) and custom indexed primers. Purified PCR libraries were amplified using Ampure XP beads, double-sided bead purification (sample to beads ratios 0.5X and 1.3X).

Whole blood was collected in a BD Vacutainer Cell Preparation Tube and processed for PBMC extraction and purification. Isolated PBMCs were frozen at -80oC in CoolCell Containers (Corning) and stored in liquid nitrogen. Frozen PBMCs were thawed in a 37°C water bath for 2-3 min. An aliquot of PBMC were stained with DAPI and counted using Countess III Automated Cell Counter. An aliquot of 50,000 cells was added to 1 ml of ATAC-RSB and spun at 1000 g for 10 minutes at 4⁰C. The supernatant was removed without disturbing the pellet. The nuclei pellet was resuspended by pipetting in 50 μL of transposition mixture (2.5μL Transposase, 25 μL 2x tagmentation buffer, 16.5 μL PBS, 0.5 μL 1% digitonin, 0.5 μL 10% Tween-20, 5 μL water) and incubated at 37°C on a thermomixer for 30 minutes with 1000 rpm shaking. The transposed DNA was purified using Qiagen MinElute Purification kits (Qiagen, 28006), and amplified using NEBnext High-Fidelity 2x PCR Master Mix (NEB, M0541L) and custom indexed primers. 90 μL SPRIselect beads (beads : DNA = 1.8) were used to purify the PCR reaction and remove excess primers and primer dimers.

#### ATAC-seq library sequencing and data processing

Pooled libraries were sequenced on an Illumina NovaSeq 6000 platform (Illumina, San Diego, CA, USA) using a paired-end 100 base-pair run configuration to a target depth of 25 million read pairs (50 million paired-end reads) per sample for muscle and to a target depth of 35 million read pairs for PBMCs. Reads were demultiplexed with bcl2fastq2 (v2.20.0) (Illumina, San Diego, CA, USA). Data was processed with the ENCODE ATAC-seq pipeline (v1.7.0) (https://github.com/ENCODE-DCC/atac-seq-pipeline). Samples from a single sex, group and exercise timepoint, e.g., endurance exercising males 30 minutes post exercise bout, were analyzed together as biological replicates in a single workflow. Briefly, adapters were trimmed with cutadapt (v2.5) and aligned to human genome hg38 with Bowtie 2 (v2.3.4.3) ^106^. Duplicate reads and reads mapping to the mitochondrial chromosome were removed. Signal files and peak calls were generated using MACS2 v2.2.4 ^107^, both from reads from each sample and pooled reads from all biological replicates. Pooled peaks were compared with the peaks called for each replicate individually using Irreproducibility Discovery Rate and thresholded to generate an optimal set of peaks. The cloud implementation of the ENCODE ATAC-seq pipeline and source code for the post-processing steps are available at https://github.com/MoTrPAC/motrpac-atac-seq-pipeline. Optimal peaks (overlap.optimal_peak.narrowPeak.bed.gz) from all workflows were concatenated, trimmed to 200 base pairs around the summit, and sorted and merged with bedtools v2.29.0 to generate a master peak list ^108^. This peak list was intersected with the filtered alignments from each sample using bedtools coverage with options -nonamecheck and -counts to generate a peak by sample matrix of raw counts. The remaining steps were applied separately on raw counts from each tissue. Peaks from non-autosomal chromosomes were removed, as well as peaks that did not have at least 10 read counts in six samples, which aided in increasing significant peak output in differential analysis. Principal component analysis (PCA) was conducted to find sample outliers to be excluded from downstream analysis using the call_pca_outliers command (found in the https://github.com/MoTrPAC/MotrpacRatTraining6mo/R package) ^109^. No PCA outliers were identified and, thus, no samples were excluded from the study. Filtered raw counts were then quantile-normalized with variancePartition::voomWithDreamWeights ^110^, which is akin to limma::voom ^111^, with the added ability to include random effects in the formula.

#### Transcriptomics

RNA Sequencing (RNA-Seq) was performed at Stanford University and the Icahn School of Medicine at Mount Sinai. Processing randomization for blood, muscle was done according to https://github.com/MoTrPAC/clinical-sample-batching. See below for adipose randomization considerations.

#### Extraction of total RNA

Tissues (∼10 mg for muscle, ∼50 mgs for adipose) were disrupted in Agencourt RNAdvance tissue lysis buffer (Beckman Coulter, Brea, CA) using a tissue ruptor (Omni International, Kennesaw, GA, #19-040E). Total RNA was extracted in a BiomekFX automation workstation according to the manufacturer’s instructions for tissue-specific extraction. Total RNA from 400 μL of blood collected in PAXgene tubes (BD Biosciences, Franklin Lakes, NJ, # 762165) was extracted using the Agencourt RNAdvance blood specific kit (Beckman Coulter). Two tissue-specific consortium reference standards were included to monitor the sample processing QC. The RNA was quantified by NanoDrop (ThermoFisher Scientific, # ND-ONE-W) and Qubit assay (ThermoFisher Scientific), and the quality was determined by either Bioanalyzer or Fragment Analyzer analysis.

#### mRNA Sequencing Library Preparation

Universal Plus mRNA-Seq kit from NuGEN/Tecan (# 9133) were used for generation of RNA-Seq libraries derived from poly(A)-selected RNA according to the manufacturer’s instructions. Universal Plus mRNA-Seq libraries contain dual (i7 and i5) 8 bp barcodes and an 8 bp unique molecular identifier (UMI), which enable deep multiplexing of NGS sequencing samples and accurate quantification of PCR duplication levels. Approximately 500ng of total RNA was used to generate the libraries for muscle, 300ng for adipose, and 250ng of total RNA was used for blood. The Universal Plus mRNA-Seq workflow consists of poly(A) RNA selection, RNA fragmentation and double-stranded cDNA generation using a mixture of random and oligo(dT) priming, end repair to generate blunt ends, adaptor ligation, strand selection, AnyDeplete workflow to remove unwanted ribosomal and globin transcripts, and PCR amplification to enrich final library species. All library preparations were performed using a Biomek i7 laboratory automation system (Beckman Coulter). Tissue-specific reference standards provided by the consortium were included with all RNA isolations to QC the RNA. RNA Sequencing, quantification, and normalization

RNA sequencing, quantification, and normalization Pooled libraries were sequenced on an Illumina NovaSeq 6000 platform (Illumina, San Diego, CA, USA) to a target depth of 40 million read pairs (80 million paired-end reads) per sample using a paired-end 100 base pair run configuration. In order to capture the 8-base UMIs, libraries were sequenced using 16 cycles for the i7 index read and 8 cycles for the i5 index read. Reads were demultiplexed with bcl2fastq2 (v2.20.0) using options --use-bases-mask Y*,I8Y*,I*,Y* --mask-short-adapter-reads 0 --minimum-trimmed-read-length 0 (Illumina, San Diego, CA, USA), and UMIs in the index FASTQ files were attached to the read FASTQ files. Adapters were trimmed with cutadapt (v1.18) ^106^, and trimmed reads shorter than 20 base pairs were removed. FastQC (v0.11.8) was used to generate pre-alignment QC metrics. STAR (v2.7.0d) ^112^ was used to index and align reads to release 38 of the Ensembl Homo sapiens (hg38) genome and Gencode (Version 29). Default parameters were used for STAR’s genomeGenerate run mode; in STAR’s alignReads run mode, SAM attributes were specified as NH HI AS NM MD nM, and reads were removed if they did not contain high-confidence collapsed splice junctions (--outFilterType BySJout). RSEM (v1.3.1) ^113^ was used to quantify transcriptome-coordinate-sorted alignments using a forward probability of 0.5 to indicate a non-strand-specific protocol. Bowtie 2 (v2.3.4.3) ^114^ was used to index and align reads to globin, rRNA, and phix sequences in order to quantify the percent of reads that mapped to these contaminants and spike-ins. UCSC’s gtfToGenePred was used to convert the hg38 gene annotation (GTF) to a refFlat file in order to run Picard CollectRnaSeqMetrics (v2.18.16) with options MINIMUM_LENGTH=50 and RRNA_FRAGMENT_PERCENTAGE=0.3. UMIs were used to accurately quantify PCR duplicates with NuGEN’s “nodup.py” script (https://github.com/tecangenomics/nudup). QC metrics from every stage of the quantification pipeline were compiled, in part with multiQC (v1.6). The openWDL-based implementation of the RNA-Seq pipeline on Google Cloud Platform is available on Github (https://github.com/MoTrPAC/motrpac-rna-seq-pipeline). Filtering of lowly expressed genes and normalization were performed separately in each tissue. RSEM gene counts were used to remove lowly expressed genes, defined as having 0.5 or fewer counts per million in at least 10% of samples. These filtered raw counts were used as input for differential analysis with the variancePartition::dream ^110^, as described in the statistical analysis methods section. To generate normalized sample-level data for downstream visualization, filtered gene counts were TMM-normalized using edgeR::calcNormFactors, followed by conversion to log counts per million with edgeR::cpm.

Principal Component Analysis and calculation of the variance explained by variables of interest were used to identify and quantify potential batch effects. Based on this analysis, processing batch (muscle and blood, see below for adipose), Clinical Site (all tissues), percentage of UMI duplication, and RNA Integrity Number (RIN) technical effects were regressed out of the TMM-normalized counts via linear regression using limma::RemoveBatchEffect function in R ^115^. A design matrix including age, sex, and a combination of group and timepoint was used during batch effect removal to avoid removing variance attributable to biological effects.

#### Adipose tissue processing considerations

In the adipose tissue, RNA extraction was performed in separate batches according to exercise modality, so the extraction batch and exercise modality were perfectly collinear. This collinearity was identified at the RNA extraction step and samples were randomized prior to construction of cDNA libraries. Ultimately, under this processing implementation, differences in gene expression attributable to exercise group can be impossible to disentangle from RNA extraction batch effects, so the batch variable was not regressed out in the technical effect stage or included as a covariate in the differential analysis, and left as an experimental limitation.

#### RNA Quality Inclusion Criteria

In the blood transcriptomic data, 79/1032 processed adult-sedentary samples had RIN values under 5. Based on established guidelines and internal quality control assessments, any samples with a RIN score below 5 were excluded from further analysis due to concerns about potential degradation artifacts. Additional visualizations and summary figures supporting this decision are available in the quality control report at: https://github.com/MoTrPAC/precovid-analyses/tree/main/QC/

### Proteomics and Phosphoproteomics

#### Study design

LC-MS/MS analysis of 379 muscle samples encompassing baseline and 3 post-intervention timepoints from all three groups (control, endurance and resistance) was performed at the Broad Institute of MIT and Harvard (BI) and Pacific Northwest National Laboratories (PNNL). Samples were split evenly across the two sites, and a total of 14 samples were processed and analyzed at both sites to serve as cross-site replicates for evaluation of reproducibility. Additionally, 46 adipose tissue samples representing baseline and 4-hours post-intervention from all three groups were analyzed at PNNL.

#### Generation of common reference

For both tissue types, a tissue-specific common reference material was generated from bulk human samples. The common reference sample for muscle consisted of bulk tissue digest from 5 individuals at 2/3 ratio of female/male. Samples were split equally between BI and PNNL, digested at each site following sample processing protocol described below, then mixed all digests from both sites and centrally aliquoted at PNNL. Common reference for adipose tissue was generated at PNNL using bulk tissue from 6 individuals representing a 4:2 ratio of female:male. 250 μg aliquots of both tissue specific common reference samples were made to be included in each multiplex (described below) and additional aliquots are stored for inclusion in future MoTrPAC phases to facilitate data integration.

#### Sample processing

Proteomics analyses were performed using clinical proteomics protocols described previously ^116,117^. Muscle and adipose samples were lysed in ice-cold, freshly-prepared lysis buffer (8 M urea (Sigma-Aldrich, St. Louis, Missouri), 50 mM Tris pH 8.0, 75 mM sodium chloride, 1 mM EDTA, 2 μg/ml Aprotinin (Sigma-Aldrich, St. Louis, Missouri), 10 μg/ml Leupeptin (Roche CustomBiotech, Indianapolis, Indiana), 1 mM PMSF in EtOH, 10 mM sodium fluoride, 1% phosphatase inhibitor cocktail 2 and 3 (Sigma-Aldrich, St. Louis, Missouri), 10 mM Sodium Butyrate, 2 μM SAHA, and 10 mM nicotinamide and protein concentration was determined by BCA assay. Protein lysate concentrations were normalized within samples of the same tissue type, and protein was reduced with 5 mM dithiothreitol (DTT, Sigma-Aldrich) for 1 hour at 37°C with shaking at 1000 rpm on a thermomixer, alkylated with iodoacetamide (IAA, Sigma-Aldrich) in the dark for 45 minutes at 25°C with shaking at 1000 rpm, followed by dilution of 1:4 with Tris-HCl, pH 8.0 prior to adding digestion enzymes. Proteins were first digested with LysC endopeptidase (Wako Chemicals) at a 1:50 enzyme:substrate ratio (2 hours, 25 °C, 850 rpm), followed by digestion with trypsin (Promega) at a 1:50 enzyme:substrate ratio (or 1:10 ratio for adipose tissue; 14 hours, 25 °C, 850 rpm). The next day formic acid was added to a final concentration of 1% to quench the reaction. Digested peptides were desalted using Sep-Pac C18 columns (Waters), concentrated in a vacuum centrifuge, and a BCA assay was used to determine final peptide concentrations. 250μg aliquots of each sample were prepared, dried down by vacuum centrifugation and stored at -80°C.

Tandem mass tag (TMT) 16-plex isobaric labeling reagent (ThermoFisher Scientific) was used for this study. Samples were randomized across the first 15 channels of TMT 16-plexes, and the last channel (134N) of each multiplex was used for a common reference that was prepared prior to starting the study (see above). Randomization of samples across the plexes within each site was done using https://github.com/MoTrPAC/clinical-sample-batching, with the goal to have all timepoints per participant in the same plex, and uniform distribution of groups (endurance, resistance, control), sex and sample collection clinical site across the plexes.

Peptide aliquots (250 μg per sample) were resuspended to a final concentration of 5 μg/μL in 200 mM HEPES, pH 8.5 for isobaric labeling. TMT reagent was added to each sample at a 1:2 peptide: TMT ratio, and labeling proceeded for 1 hour at 25°C with shaking at 400 rpm. The labeling reaction was diluted to a peptide concentration of 2 µg/µL using 62.5 μL of 200 mM HEPES and 20% ACN. 3 μL was removed from each sample to quantify labeling efficiency and mixing ratio. After labeling QC analysis, reactions were quenched with 5% hydroxylamine and samples within each multiplex were combined and desalted with Sep-Pac C18 columns (Waters).

Combined TMT multiplexed samples were then fractionated using high pH reversed phase chromatography on a 4.6mm ID x 250mm length Zorbax 300 Extend-C18 column (Agilent) with 5% ammonium formate/2% Acetonitrile as solvent A and 5% ammonium formate/90% acetonitrile as solvent B. Samples were fractionated with 96min separation gradient at flow rate of 1mL/min and fractions were collected at each minute onto a 96-well plate. Fractions are then concatenated into 24 fractions with the following scheme: fraction 1 = A1+C1+E1+G1, fraction 2 = A2+C2+E2+G2, fraction 3 = A3+C3+E3+G3, all the way to fraction 24 = B12+D12+F12+H12 following the same scheme. 5% of each fraction was removed for global proteome analysis, and the remaining 95% was further concatenated to 12 fractions for phosphopeptide enrichment using immobilized metal affinity chromatography (IMAC).

Phosphopeptide enrichment was performed through immobilized metal affinity chromatography (IMAC) using Fe 3+ -NTA-agarose beads, freshly prepared from Ni-NTA-agarose beads (Qiagen, Hilden, Germany) by sequential incubation in 100 mM EDTA to strip nickel, washing with HPLC water, and incubation in 10 mM iron (III) chloride). Peptide fractions were resuspended to 0.5 μg/uL in 80% ACN + 0.1% TFA and incubated with beads for 30 minutes in a thermomixer set to 1000 rpm at room temperature. After 30 minutes, beads were spun down (1 minute, 1000 rcf) and supernatant was removed and saved as flow-through for subsequent enrichments. Phosphopeptides were eluted off IMAC beads in 3x 75 μL of agarose bead elution buffer (500 mM K2HPO4, pH 7.0), desalted using C18 stage tips, eluted with 50% ACN, and lyophilized. Samples were then reconstituted in 3% ACN / 0.1% FA for LC-MS/MS analysis (9 μL reconstitution / 4 μL injection at the BI; 12 μL reconstitution / 5 μL injection at PNNL).

#### Data acquisition

Broad Institute: Both proteome and phosphoproteome samples were analyzed on 75um ID Picofrit columns packed in-house with ReproSil-Pur 120 Å, C18-AQ, 1.9 µm beads to the length of 20-24cm. Online separation was performed on Easy nLC 1200 systems (ThermoFisher Scientific) with solvent A of 0.1% formic acid/3% acetonitrile and solvent B of 0.1% formic acid/90% acetonitrile, flow rate of 200nL/min and the following gradient: 2-6% B in 1min, 6-20% B in 52min, 20-35% B in 32min, 35-60% B in 9min, 60-90% B in 1min, followed by a 5min hold at 90%, and 9min hold at 50%. Proteome fractions were analyzed on a Q Exactive Plus mass spectrometer (ThrmoFisher Scientific) with MS1 scan across the 300-1800 m/z range at 70,000 resolution, AGC target of 3×106 and maximum injection time of 5ms. MS2 scans of most abundant 12 ions were performed at 35,000 resolution with AGC target of 1×105 and maximum injection time of 120ms, isolation window of 0.7m/z and normalized collision energy of 27. Phosphoproteome fractions were analyzed on Q Exactive HFX MS system (ThermoFisher Scientific) with the following parameters: MS1 scan across 350-1800 m/z mass range at 60,000 resolution, AGC target of 3×106 and maximum injection time of 10ms. 20 most abundant ions are fragmented in MS2 scans with AGC of 1×105, maximum injection time of 105ms, isolation width of 0.7 m/z and NCE of 29. In both methods dynamic exclusion was set to 45sec.

PNNL: For mass spectrometry analysis of the global proteome of muscle samples, online separation was performed using a nanoAcquity M-Class UHPLC system (Waters) and a 25 cm x 75 μm i.d. picofrit column packed in-house with C18 silica (1.7 μm UPLC BEH particles, Waters Acquity) with solvent A of 0.1% formic acid/3% acetonitrile and solvent B of 0.1% formic acid/90% acetonitrile, flow rate of 200nL/min and the following gradient: 1% B for 8min, 8-20% B in 90min, 20-35% B in 13min, 35-75% B in 5min, 75-95% B in 3min, followed by a 6min hold at 90%, and 9min hold at 50%. Proteome fractions were analyzed on a Q Exactive Plus mass spectrometer (ThermoFisher Scientific) with MS1 scan across the 300-1800 m/z range at 60,000 resolution, AGC target of 3×106 and maximum injection time of 20ms. MS2 scans of most abundant 12 ions were performed at 30,000 resolution with AGC target of 1×105 and maximum injection time of 100ms, isolation window of 0.7m/z and normalized collision energy of 30.

For mass spectrometry analysis of the global proteome of adipose samples, online separation was performed using a Dionex Ultimate 3000 UHPLC system (ThermoFisher) and a 25 cm x 75 μm i.d. picofrit column packed in-house with C18 silica (1.7 μm UPLC BEH particles, Waters Acquity) with solvent A of 0.1% formic acid/3% acetonitrile and solvent B of 0.1% formic acid/90% acetonitrile, flow rate of 200nL/min and the following gradient: 1-8% B in 10min, 8-25% B in 90min, 25-35% B in 10min, 35-75% B in 5min, and 75-5% in 3min. Proteome fractions were analyzed on a Q Exactive HF-X Plus mass spectrometer (ThermoFisher Scientific) with MS1 scan across the 300-1800 m/z range at 60,000 resolution, AGC target of 3×106 and maximum injection time of 20ms. MS2 scans of most abundant 12 ions were performed at 30,000 resolution with AGC target of 1×105 and maximum injection time of 100ms, isolation window of 0.7m/z and normalized collision energy of 30.

For mass spectrometry analysis of the phosphoproteome of both sample types, online separation was performed using a Dionex Ultimate 3000 UHPLC system (ThermoFisher) and a 25 cm x 75 μm i.d. picofrit column packed in-house with C18 silica (1.7 μm UPLC BEH particles, Waters Acquity) with solvent A of 0.1% formic acid/3% acetonitrile and solvent B of 0.1% formic acid/90% acetonitrile, flow rate of 200nL/min and the following gradient: 1-8% B in 10min, 8-25% B in 90min, 25-35% B in 10min, 35-75% B in 5min, and 75-5% in 3min. Phosphoproteome fractions were analyzed on a Q Exactive HF-X Plus mass spectrometer (ThermoFisher Scientific) with MS1 scan across the 300-1800 m/z range at 60,000 resolution, AGC target of 3×106 and maximum injection time of 20ms. MS2 scans of most abundant 12 ions were performed at 45,000 resolution with AGC target of 1×105 and maximum injection time of 100ms, isolation window of 0.7m/z and normalized collision energy of 30.

#### Data searching

Proteome and phosphoproteome data from both BI and PNNL were searched against a composite protein database at the Bioinformatics Center (BIC) using the MSGF+ cloud-based pipeline previously described ^118^. This database comprised UniProt canonical sequences (downloaded 2022-09-13; 20383 sequences), UniProt human protein isoforms (downloaded 2022-09-13; 21982 sequences), and common contaminants (261 sequences), resulting in 42,626 sequences.

#### QC/Filtering/Normalization

The log2 Reporter ion intensity (RII) ratios to the common reference were used as quantitative values for all proteomics features (proteins and phosphosites). Datasets were filtered to remove features identified from contaminant proteins and decoy sequences. Datasets were visually evaluated for sample outliers by looking at top principal components, examining median feature abundance and distributions of RII ratio values across samples, and by quantifying the number of feature identifications within each sample. No outliers were detected in the muscle tissue dataset. In the adipose datasets (proteome and phosphoproteome), four samples were flagged as outliers based on inspection of the top principal components and evaluation of the raw quantitative results (median feature abundance <-1.0). These outlier samples were removed from the dataset, and the list of outlier samples can be found in Supplementary Table S1. The Log2 RII ratio values were normalized within each sample by median centering to zero. Principal Component Analysis and calculation of the variance explained by variables of interest were used to identify and quantify potential batch effects. Based on this analysis, TMT plex (muscle and adipose), Clinical Site (muscle and adipose), and Chemical Analysis Site (muscle only, where samples were analyzed at both BI and PNNL) batch effects were removed using Linear Models Implemented in the limma::RemoveBatchEffect() function in R ^115^. A design matrix including age, sex, and group_timepoint was used during batch effect removal in order to preserve the effect of all variables included in later statistical analysis. Correlations between technical replicates analyzed within and across CAS (where applicable) were calculated to evaluate intra-and inter-site reproducibility; the data from technical replicates were then averaged for downstream analysis. Finally, features with quantification in less of 30% of all samples were removed. For specific details of the process, see available code.

### Statistical analysis for gene level RNA expression, chromatin accessibility and phosphoprotein expression

This section briefly describes the statistical analysis of the non-splicing omics, which is the focus of the Multi-omic Landscape. These include gene-level RNA expression, chromatin accessibility, and protein and phosphoprotein expression. These omics data were modeled similarly because they are non-negative quantity measures. The full implementation of the statistical models for these omics can be found in the R Package

MotrpacHumanPreSuspensionData. (https://github.com/MoTrPAC/MotrpacHumanPreSuspensionData). In this manuscript, statistical analysis results on these omics were directly obtained from this R package. The statistical analysis of alternative splicing is described in detail in the next section “Statistical analysis of alternative splicing”, because alternative splicing is quantified by PSI values with a different statistical distribution than the omics in this section.

#### Differential analysis

To model the effects of both exercise modality and time, relative to non exercising control, each measured molecular feature was treated as an outcome in a linear mixed effects model accounting for fixed effects of exercise group (RE, EE, or CON), timepoint, as well as demographic and technical covariates (see covariate selection below for more info). Participant identification was treated as a random effect. For each molecular feature, a cell-means model is fit to estimate the mean of each exercise group-timepoint combination, and all hypothesis tests are done comparing the means of the fixed effect group-timepoint combinations. In order to model the effects of exercise against non-exercising control, a difference-in-changes model was used which compared the change from pre-exercise to a during or post-exercise timepoint in one of the two exercise groups to the same change in control. This effect is sometimes referred to as a “delta-delta” or “difference-in-differences” model. See Multi-omic Landscape for detailed description of model selection, simulation studies and missing data handling.

#### Covariate selection

Covariates were selected through a combination of a priori knowledge about factors influencing molecular levels as well through empiric screening. Factors were considered for model inclusion by correlating the principal components of each tissue-ome feature set to demographic and technical factors using variancePartition::canCorPairs ^110^. Visualizations and computational analysis that describe this process for the decisions for covariate selection can be found at: https://github.com/MoTrPAC/precovid-analyses/tree/main/QC. Ultimately, the following fixed effect covariates were included in the models for every omic platform: group and timepoint in a cell means model, clinical site, age, sex, and BMI. ‘Participant id’ was included as a random effect in every model. Each ome then had ome-specific covariates selected as described in prior sections. See Multi-omic Landscape on a discussion of including race, ethnicity and genetic ancestry into the model.

#### Specific omic-level statistical considerations

Models for all omes were fit using ‘variancePartition::dream’.129 For the transcriptomics and ATAC-seq, which are measured as numbers of counts, the mean-variance relationship was measured using ‘variancePartition::voomWithDreamWeights’ as previously described ^110^. For all other omes the normalized values were used directly as input to the statistical model.

#### Significance thresholds

For each of the above contrasts, p-values were adjusted for multiple comparisons for each unique contrast-group-tissue-ome-timepoint combination separately using the Benjamini-Hochberg method to control False Discovery Rate (FDR) ^119^. Features were considered significant at a FDR of 0.05 unless otherwise stated.

#### Human feature to gene mapping

The feature-to-gene map links each feature tested in differential analysis to a gene, using Ensembl version 105 (mapped to GENCODE 39) ^120^ as the gene identifier source. Proteomics feature IDs (UniProt IDs) were mapped to gene symbols and Entrez IDs using UniProt’s mapping files ^121^. Epigenomics features were mapped to the nearest gene using the ChIPseeker::annotatePeak() ^122,123^ function with Homo sapiens Ensembl release 105 gene annotations. Gene symbols, Entrez IDs, and Ensembl IDs were assigned to features using biomaRt version 2.58.2 (Bioconductor 3.18) ^124–126^.

### Statistical analysis for alternative splicing

#### Alternative splicing quantification from RNA-seq

A uniform pipeline was employed to process all RNA-seq fastq files. In particular, STAR (v2.7.4)34 was used to align the reads to hg38 genome build with Gencode v30 index. To quantify gene expression levels, kallisto (v0.46.0)36 was used to pseudo-align RNA-seq reads to Gencode v38 transcripts. Throughout this study, Gencode v30 genome annotation was used as the reference gene annotations wherever applicable.

To reduce potential counting bias, two distinct approaches leveraging different aspects of RNA-seq reads to quantify Percent Spliced In (PSI) were employed and combined. Using genome read alignment generated by STAR as input, the junction read counts for alternative splicing events were counted by DARTS/rMATS-turbo (rMATS) ^41,42^. Using the transcript quantifications generated by kallisto as input, the ratio between longer and shorter isoforms were computed by SUPPA2 (SUPPA) ^43^. We analyzed four basic types of alternative splicing events, i.e., skipped exons, alternative 5′ splice sites, alternative 3′ splice sites, and retained introns.

#### AS events filtering

AS events characterized by rMATS and SUPPA were filtered based on the abundance of reads (for rMATS) or transcripts (for SUPPA) associated with each AS event. Specifically, we only keep AS events for which the corresponding abundance values were no less than a cutoff in at least 20% of the samples, unless this AS event was detected by both SUPPA and rMATS. The cut off value is 5 reads for rMATS and 0.5 in TPM for SUPPA.

#### Combined AS events from rMATS and SUPPA

Because rMATS and SUPPA characterize and quantify AS events in distinct ways, we combined the analysis results from both methods for a more comprehensive coverage of all possible AS events in the data. Specifically, when an AS event was only characterized by one method, we used the PSI value from that method for this AS event in all analyses. When an AS event was characterized by both rMATS and SUPPA, with different PSI values, both PSI values were tested in the downstream analyses, including DAS events discovery, RBP analysis and splice regulatory circuitry identification. The AS event was reported as significant if one of the two PSI values returned significant result, and the result was also annotated which method (rMATS or SUPPA) this result was based on.

#### Differential alternative splicing analysis

Differential AS (DAS) events were identified using a linear mixed model. Specifically, for the samples of each exercise group G (EE, RE or Control) and each time point T during or post exercise, we concatenated them with the corresponding pre-exercise samples of the same group G, and used the following mixed model on the PSI values of each AS event:

logit (PSI) ∼ Timepoint + Batch + BMI + calculatedAge + codedsiteid + pct_umi_dup + RIN + Sex + (1 | pid)

Specifically, the logit transform of PSI value was done using the “car::logit” function in R with “adjust = 0.001”. The “lmerTest::lmer” function was used for building the model and the t-test p value for the coefficient “Timepoint” was obtained from the “summary” function. This p value is the significance p value of this AS event at time point T in Group G. P values were adjusted by FDR within the same time point and group.

To leverage the control group for the removal of circadian effects, a significant DAS event for time point T in the EE or RE group is defined as a DAS event that satisfies:

1. FDR < 0.05 in the EE or RE group, AND;
2. Satisfy one of the following:

1.1 FDR > 0.05 in the Control group, OR;
1.2 FDR < 0.05 in the Control group but the direction of change in the Control group is different from the EE or RE group.

#### Protein-coding sequence annotation

To identify AS events that directly changes CDS, the alternative exon region of each DAS event were annotated as whether they overlap with annotated CDS using the “EnsDb.Hsapiens.v86” package for the annotation and the “ensembldb::genomeToProtein” function for mapping genome coordinates to CDS annotations.

To identify AS events that are associated with a likely change of CDS, we used the SUPPA characterization of AS events. For each AS event, there was a set of RNA isoforms associated with the inclusion form of this AS event, and a distinct set of RNA isoforms associated with the exclusion form of this AS event. We obtained the annotated protein sequence of these two sets of RNA isoforms using the “ensembldb::transcripts” function. If the protein sequences showed no overlap between these two sets, this AS event was annotated as a likely change of CDS, unless it has been annotated as a direct CDS change as described above.

#### Functional annotation enrichment

We used the “ClusterProfiler::enrichGO” ^127,128^ function to find the Gene Ontology (GO) terms enriched in the genes with DAS events at each time point and groups. We used the “ClusterProfiler::simplify” function to remove redundant GO terms from the results.

### Integrative analysis of splicing regulation

#### WGCNA

Weighted Gene Co-expression Network Analysis (WGCNA; ^52^) was performed to identify modules of co-expressed genes in baseline (pre-exercise) skeletal muscle transcriptome data from 175 participants. Following QC and normalization steps, the gene expression data were adjusted for age, sex, and BMI using linear regression, and the resulting residuals from these models were used as input for network analysis. A signed WGCNA network was constructed by choosing a soft-thresholding power β to achieve scale-free topology (R² > 0.85). This process resulted in the identification of 17 co-expressing modules. Functional annotation of these modules was achieved by performing Gene Ontology (GO) enrichment analysis for each module using the clusterProfiler package ^127,128^ in R. Significance of GO terms was defined by Benjamini–Hochberg adjusted p-value < 0.05. Redundancy in the significantly enriched GO terms was addressed by performing hierarchical clustering based on kappa similarity metric, as implemented in Metascape ^129^. The most significant GO term from each cluster was then used to assign a module’s predominant biological function.

#### Spliceosome analysis

The list of components of the spliceosome machinery was downloaded from the KEGG database (ko03040 Spliceosome).

#### RBP target enrichment analysis

eCLIP data containing the genomic coordinates of the binding sites of human RBPs were downloaded from ENCODE by filtering for eCLIP datasets from the functional genomics experiment search. Track .bed files belonging to the same RBP were merged. For each RBP, the target AS events were identified by the overlap between the binding site of this RBP and a +/-100 bp window surrounding the splice site of each AS event. Enrichment of RBP binding sites in the DAS events at each timepoint and each exercise group were calculated using a hypergeometrical test with all characterized AS events as the background.

The correlation between RBP phosphorylation log2 fold changes and PSI value differences of AS events were calculated across participants where both protein phosphorylation and RNAseq were available. For each time point post exercise and each exercise modality, the within-participant log2 fold changes of phosphorylation values of all measured phosphorylation sites of RBPs were calculated by comparing the time point with pre-exercise. The changes of PSI values of all DAS events were calculated for the same participants. Pearson correlation and p values of t-test were calculated between the phosphorylation changes and the PSI changes across all participants in all time points and exercise modalities using the “psych::corr.test” function. P values were adjusted by FDR and RBP-AS event pairs with FDR < 0.05 were reported as significant.

#### Splice regulatory circuit analysis

We adapted the MAGICAL framework ^63^, which was originally designed to extract regulatory circuits for gene expression, to identify splice regulatory circuits (SRC) for the DAS events regulated by exercise. Specifically, we used all samples for which both ATACseq and RNAseq were available, and used the PSI values of AS events instead of the gene expression values in the original MAGICAL framework. Additionally, because the original MAGICAL framework was designed to run on single cell data and use the cell type pseudobulk chromatin accessibility and gene expression values, we substitute the pseudobulk calculation directly with the bulk level chromatin accessibility and PSI values. The chromatin peak to TF mapping was obtained using the r function “motifmatchr::matchMotifs” and human TF motifs from JASPAR.

We focus on finding the regulatory circuits for DAS events that were significant in at least one time point in EE or RE. For the candidate chromatin peaks, we include chromatin peaks with differential accessibility in at least one time point in EE or RE with un-adjusted p value < 0.005, and locating within a genomic window of +/- 200kb around the first splice site of at least one of the selected DAS event. The less stringent cutoff for the differential peaks was due to the lower strength of signal in the ATACseq assay, and also because the analysis focuses on identifying cross-individual correlation patterns and doesn’t rely on a stringent differential cutoff. We only keep SRCs with MAGICAL-calculated posterior looping probability >= 0.9 and TF binding probability >= 0.9. When plotting the correlation between assays for an example SRC, the pearson correlation and p values of t-test were calculated using the “psych::corr.test” function and the p value was un-adjusted because the correlation p value was not used by MAGICAL for significance calling.

## Acknowledgements

The MoTrPAC Study is supported by NIH grants U24OD026629 (Bioinformatics Center), U24DK112349, U24DK112342, U24DK112340, U24DK112341, U24DK112326, U24DK112331, U24DK112348 (Chemical Analysis Sites), U01AR071133, U01AR071130, U01AR071124, U01AR071128, U01AR071150, U01AR071160, U01AR071158 (Clinical Centers), U24AR071113 (Consortium Coordinating Center), U01AG055133, U01AG055137 and U01AG055135 (PASS/Animal Sites).

Additional grant funding: J.M.R: K23 HL150327, R03OD038387; P.R: K23HL177335-01; D.H.K: K23HL164980, 23CDA1040581; B.R: 5U01AR071150, 2U01AR071130; E.R: P30DK072476; E. K.:R01AG066474; K.C.B:1K01HL177266-01A1; K.M.:R01AG089069; M.R:NIH 5 T32 GM135066; N.M and S.E: P30AG094848; Z.C:NIH R00 HL159241; R.G: R01NR019628, R01DK081572, 21CVD01 (Leducq Foundation), R01HL133870; M.E.L, S.M, and M.T.W: The Wu Tsai Human Performance Alliance at Stanford and th e Joe and Clara Tsai

Foundation. T.C. and E.V: P30AG044271; R.S.R: I01BX003271-05A1, 1IK6BX007133-01; M.C.: NSF 2238125, NIH R01 HG009299, NIH R01 EY030546.

## Author Information

### Authorship affiliations

**Aging and Metabolism Research Program, Oklahoma Medical Research Foundation, Oklahoma City, OK**

Sue C. Bodine

**Barshop Institute for Longevity and Aging Studies, University of Texas Health Science Center at San Antonio, San Antonio, TX**

Tiffany M. Cortes, Elena Volpi

**Biological Sciences Division, Pacific Northwest National Laboratory, Richland, WA**

Joshua N. Adkins, Isaac K. Attah, Chelsea M. Hutchinson-Bunch, Damon T. Leach, Gina M. Many, Matthew E. Monroe, Ronald J. Moore, Vladislav A. Petyuk, Wei-Jun Qian, Tyler J. Sagendorf, James A. Sanford

**Biomedical Data Science Department, Stanford University, Stanford, CA**

Trevor Hastie, Robert Tibshirani

**Birmingham/Atlanta VA GRECC, Birmingham VA Medical Center, Birmingham, AL**

Thomas W. Buford

**Blavatnik School of Computer Science and AI, Tel Aviv University, Tel Aviv,Israel**

David Amar

**BRCF Metabolomics Core, University of Michigan, Ann Arbor, MI**

Charles F. Burant, Charles R. Evans, Maureen T. Kachman, Abraham Raskind, Alexander Raskind, Tanu Soni, Angela Wiggins

**Broad Institute of MIT and Harvard, Cambridge, MA**

Robert E. Gerszten

**CardioVascular Institute, Beth Israel Deaconess Medical Center, Boston, MA**

Robert E. Gerszten, Prashant Rao, Jeremy M. Robbins

**Cardiovascular Institute, Stanford School of Medicine, Stanford, CA**

Michael P. Snyder

**Center for Translational Geroscience, Cedars-Sinai Health Sciences University, Los Angeles, CA**

Sara E. Espinoza

**Computational and Systems Biology, University of Pittsburgh, Pittsburgh, PA**

Maria Chikina

**Department of Biochemistry, Emory University School of Medicine, Atlanta, GA**

Eric A. Ortlund

**Department of Biomedical Data Science, Stanford University, Stanford, CA**

Trevor Hastie, Stephen B. Montgomery, Robert Tibshirani

**Department of Biomedical Informatics, University of Colorado Anschutz, Aurora, CO**

Kristen J. Sutton

**Department of Biostatistics and Data Science, Wake Forest University School of Medicine, Winston-Salem, NC**

Catherine Gervais, Fang-Chi Hsu, Byron C. Jaeger, Michael E. Miller, Joseph Rigdon, Courtney G. Simmons, Cynthia L. Stowe

**Department of Cell, Developmental and Integrative Biology, University of Alabama at Birmingham, Birmingham, AL**

Anna Thalacker-Mercer

**Department of Cellular & Integrative Physiology, University of Texas Health Science Center at San Antonio, San Antonio, TX**

Blake B. Rasmussen

**Department of Comparative Biosciences, University of Illinois at Urbana-Champaign, Champaign, IL**

Minghui Lu

**Department of Computational Medicine and Bioinformatics, University of Michigan, Ann Arbor, MI**

Gayatri Iyer

**Department of Genetics and Genomic Sciences, Icahn School of Medicine at Mount Sinai, New York, NY**

Martin J. Walsh

**Department of Genetics, Stanford University, Stanford, CA**

Euan A. Ashley, Clarisa Chavez Martinez, Christopher A. Jin, Nikhil Milind, Stephen B. Montgomery, Michael P. Snyder, Bingqing Zhao

**Department of Internal Medicine, Division of Physical Activity and Weight Management, University of Kansas Medical Center, Kansas City, KS**

John M. Jakicic, Renee J. Rogers

**Department of Internal Medicine, University of Michigan, Ann Arbor, MI**

Charles F. Burant, Charles R. Evans, Johanna Y. Fleischman, Alexander Raskind

**Department of Kinesiology, East Carolina University, Greenville, NC**

Nicholas T. Broskey, Joseph A. Houmard

**Department of Kinesiology, Nutrition, and Health, Miami University, Oxford, OH**

Alex Claiborne

**Department of Medicine-Division of Gastroenterology and Hepatology, University of Missouri, Columbia, MO**

R. Scott Rector

**Department of Medicine, Cedars-Sinai Medical Center, Los Angeles, CA**

Sara E. Espinoza, Nicolas Musi

**Department of Medicine, Division of Endocrinology, University of Pittsburgh, Pittsburgh, PA**

Erin E. Kershaw

**Department of Medicine, University of Alabama at Birmingham, Birmingham, AL**

Thomas W. Buford

**Department of Molecular and Integrative Physiology, University of Michigan, Ann Arbor, MI**

Charles F. Burant

**Department of Neurology, Icahn School of Medicine at Mount Sinai, New York, NY**

Mary Anne S. Amper, Yongchao Ge, Venugopalan D. Nair, Hanna Pincas, Stas Rirak, Stuart C. Sealfon, Gregory R. Smith, Mital Vasoya, Xuechen Yu, Elena Zaslavsky, Zidong Zhang

**Department of Nutrition and Exercise Physiology, University of Missouri, Columbia, MO**

R. Scott Rector

**Department of Orthopaedics, Atrium Health Wake Forest Baptist, Wake Forest University School of Medicine, Winston-Salem, NC**

David Popoli

**Department of Pathology and Laboratory Medicine, University of Vermont, Burlington, VT**

Sandra T. May, Russell Tracy

**Department of Pathology, Stanford University, Stanford, CA**

Euan A. Ashley, Stephen B. Montgomery, Kevin S. Smith, Nikolai G. Vetr, Yilin Xie

**Department of Pharmacological Sciences, Icahn School of Medicine at Mount Sinai, New York, NY**

Yifei Sun, Martin J. Walsh

**Department of Pharmacology & Cancer Biology and Department of Medicine, Duke University Medical Center, Durham, NC**

Olga Ilkayeva, Christopher B. Newgard

**Department of Physical Medicine and Rehabilitation, Atrium Health Wake Forest Baptist, Wake Forest University School of Medicine, Winston-Salem, NC**

David Popoli

**Department of Physical Therapy, School of Health Professions, Concordia University of Wisconsin, Mequon, WI**

Kevin J. Gries

**Department of Population Health Sciences, Duke University School of Medicine, Durham, NC**

Katherine A. Collins-Bennett

**Dept of Biomolecular Chemistry, University of Wisconsin, Madison**

Margaret Robinson

**Diabetes and Aging Center, Cedars-Sinai Health Sciences University, Los Angeles, CA**

Sara E. Espinoza

**Division of Cardiology, Department of Medicine, Duke University, Durham, NC**

William E. Kraus

**Division of Cardiovascular Medicine, Beth Israel Deaconess Medical Center, Boston, MA**

Jacob L. Barber, Robert E. Gerszten, Prashant Rao, Jeremy M. Robbins

**Division of Cardiovascular Medicine, Department of Medicine, Stanford University, Stanford, CA**

Euan A. Ashley, David Jimenez-Morales, Daniel H. Katz, Malene E. Lindholm, Samuel Montalvo, Matthew T. Wheeler, Jay Yu, Jimmy Zhen

**Division of Computational Medicine, Stanford University, Stanford, CA**

Daniel H. Katz

**Division of Diabetes, Endocrinology, and Metabolic Diseases, National Institute of Diabetes and Digestive and Kidney Diseases, National Institutes of Health**

Ashley Y. Xia

**Division of Endocrinology, Duke University Medical Center, Durham, NC**

Hiba Abou Assi

**Division of Endocrinology, Metabolism, and Diabetes, University of Colorado School of Medicine, Anschutz Medical Campus, Aurora, CO**

Bryan C. Bergman, Daniel H. Bessesen, Edward L. Melanson

**Division of Endocrinology, University of Pittsburgh, Pittsburgh, PA**

Maja Stefanovic-Racic

**Division of Geriatric Medicine, Department of Medicine, University of Colorado Anschutz Medical Campus, Aurora, CO**

Zachary S. Clayton, Wendy M. Kohrt, Edward L. Melanson, Kerrie L. Moreau

**Division of Rheumatology & Immunology, Duke Molecular Physiology Institute, Duke University, Durham, NC**

Kim M. Huffman

**Duke Molecular Physiology Institute, Duke University, Durham, NC**

Brian J. Andonian, Katherine A. Collins-Bennett, Olga Ilkayeva, Johanna L. Johnson, William E. Kraus, Christopher B. Newgard

**East Carolina Diabetes and Obesity Institute, East Carolina University, Greenville, NC**

Nicholas T. Broskey, Joseph A. Houmard

**Emory Integrated Lipidomics and Metabolomics Core, Emory University School of Medicine, Atlanta, GA**

Zhenxin Hou, Eric A. Ortlund

**Environmental and Molecular Sciences Division, Pacific Northwest National Laboratory, Richland, WA**

Marina A. Gritsenko, Paul D. Piehowski

**Gray School of Medical Sciences, Tel Aviv University, Tel Aviv, Israel**

David Amar

**Harvard Medical School, Boston, MA**

Robert E. Gerszten, Prashant Rao, Jeremy M. Robbins

**Health and Exercise Science, Wake Forest University, Winston-Salem, NC**

W. Jack Rejeski

**Human Performance Laboratory, Ball State University, Muncie, IN**

Alicia Belangee, Gerard A. Boyd, Anna R. Brandt, Toby L. Chambers, Clarisa Chavez Martinez, Matthew Douglass, Will A. Fountain, Aaron H. Gouw, Kevin J. Gries, Ryan P. Hughes, Dillon J. Kuszmaul, Bridget Lester, Colleen E. Lynch, Cristhian Montenegro, Masatoshi Naruse, Ulrika Raue, Ethan Robbins, Kaitlyn R. Rogers, Chad M. Skiles, Andrew M. Stroh, Scott Trappe, Todd A. Trappe, Caroline S. Vincenty, Gilhyeon Yoon

**Human Performance Laboratory, East Carolina University, Greenville, NC**

Nicholas T. Broskey, Joseph A. Houmard

**Icahn School of Medicine at Mount Sinai, New York, NY**

Minghui Lu, Nada Marjanovic, German Nudelman, Nora-Lovette Okwara, Alexandria Vornholt

**Larner College of Medicine at The University of Vermont, Burlington, VT**

Jessica L. Rooney

**Metabolomics Platform, Broad Institute of MIT and Harvard, Cambridge, MA**

Clary B. Clish

**National Security Directorate, Pacific Northwest National Laboratory, Richland, WA**

Joshua R. Hansen

**Neuromuscular Research Laboratory/Warrior Human Performance Research Center, Department of Sports Medicine and Nutrition, University of Pittsburgh, Pittsburgh, PA**

Bradley C Nindl

**NextGen Precision Health, University of Missouri, Columbia, MO**

R. Scott Rector

**Oregon Health & Science University, Department of Biomedical Engineering, Portland, OR**

Joshua N. Adkins

**Pediatric Exercise and Genomics Research Center, Department of Pediatrics, School of Medicine, University of California Irvine, CA**

Dan M. Cooper, Fadia Haddad, Shlomit Radom-Aizik

**Pennington Biomedical Research Center, LSU System, Baton Rouge, LA**

Tuomo Rankinen, Eric Ravussin

**Perisphere Real World Evidence, Austin, TX**

Byron C. Jaeger

**Petit Institute of Bioengineering and Bioscience, Georgia Institute of Technology, Atlanta, GA**

Facundo M. Fernandez

**Preventive Medicine, Pennington Biomedical Research Center, Baton Rouge, LA**

Neil M. Johannsen

**Proteomics Platform, Broad Institute of MIT and Harvard, Cambridge, MA**

Kevin Bonanno, Steven A. Carr, Natalie M. Clark, Patrick Hart, Pierre M. Jean-Beltran, Hasmik Keshishian, D. R. Mani, Margaret Robinson

**Research Service, Harry S Truman Memorial Veterans Medical Center, Columbia, MO**

R. Scott Rector

**San Antonio GRECC, South Texas Veterans Health Care System, San Antonio, TX**

Tiffany M. Cortes, Elena Volpi

**School of Chemistry and Biochemistry, Georgia Institute of Technology, Atlanta, GA**

Facundo M. Fernandez, David A. Gaul

**Sylvan Adams Sports Science Institute, Tel Aviv University, Tel Aviv, Israel**

David Amar

**The Mount Sinai Center for RNA Biology and Medicine, Icahn School of Medicine at Mount Sinai, New York, NY**

Martin J. Walsh

**Translational Research Institute, AdventHealth, Orlando, FL**

Cheehoon Ahn, Paul M. Coen, Bret H. Goodpaster, Lauren M. Sparks, Katie L. Whytock

**UAB School of Public Health Department of Biostatistics: Institute for Human Machine and Cognition**

Gary R. Cutter

**University of Colorado Anschutz College of Nursing, Aurora, CO**

Catherine M. Jankowski

**University of Colorado Anschutz Medical Campus, Aurora, CO**

Irene E. Schauer, Robert S. Schwartz

**University of Pittsburgh, Pittsburgh, PA**

Daniel E. Forman, Renee J. Rogers

**Veterans Administration Eastern Colorado Health Care System, Geriatric Research, Education, and Clinical Center, CO**

Kerrie L. Moreau

**Wu Tsai Human Performance Alliance, Stanford, CA**

Samuel Montalvo

### Authorship information

**Writing Group Members:** Zidong Zhang, German Nudelman, Hanna Pincas, Gayatri Iyer, Gregory Smith, Hasmik Keshishian, Christopher A. Jin, Scott Trappe, Daniel Katz, Charles Burant, Venugopalan D. Nair, Elena Zaslavsky, Stuart C. Sealfon

#### MoTrPAC Study Group members

**Bioinformatics Center:** David Amar, Trevor Hastie, David Jimenez-Morales, Daniel H. Katz, Malene E. Lindholm, Samuel Montalvo, Robert Tibshirani, Jay Yu, Jimmy Zhen, Euan A. Ashley, Matthew T. Wheeler

**Biospecimens Repository:** Sandra T. May, Jessica L. Rooney, Russell Tracy

**Data Management, Analysis, and Quality Control Center:** Catherine Gervais, Fang-Chi Hsu, Byron C. Jaeger, David Popoli, Joseph Rigdon, Courtney G. Simmons, Cynthia L. Stowe, Michael E. Miller

**Exercise Intervention Core:** W. Jack Rejeski

**NIH:** Ashley Y. Xia

**Preclinical Animal Study Sites:** Sue C. Bodine, R. Scott Rector

**Chemical Analysis Sites:** Hiba Abou Assi, Mary Anne S. Amper, Brian J. Andonian, Isaac K. Attah, Jacob L. Barber, Kevin Bonanno, Clarisa Chavez Martinez, Natalie M. Clark, Johanna Y. Fleischman, David A. Gaul, Yongchao Ge, Marina A. Gritsenko, Joshua R. Hansen, Patrick Hart, Zhenxin Hou, Chelsea M. Hutchinson-Bunch, Olga Ilkayeva, Gayatri Iyer, Pierre M. Jean-Beltran, Christopher A. Jin, Maureen T. Kachman, Hasmik Keshishian, Damon T. Leach, Minghui Lu, D. R. Mani, Gina M. Many, Nada Marjanovic, Nikhil Milind, Matthew E. Monroe, Ronald J. Moore, Venugopalan D. Nair, German Nudelman, Nora-Lovette Okwara, Vladislav A. Petyuk, Paul D. Piehowski, Hanna Pincas, Wei-Jun Qian, Prashant Rao, Abraham Raskind, Alexander Raskind, Stas Rirak, Jeremy M. Robbins, Margaret Robinson, Tyler J. Sagendorf, James A. Sanford, Gregory R. Smith, Kevin S. Smith, Yifei Sun, Mital Vasoya, Nikolai G. Vetr, Alexandria Vornholt, Yilin Xie, Xuechen Yu, Elena Zaslavsky, Zidong Zhang, Bingqing Zhao, Joshua N. Adkins, Charles F. Burant, Steven A. Carr, Clary B. Clish, Facundo M. Fernandez, Robert E. Gerszten, Stephen B. Montgomery, Christopher B. Newgard, Eric A. Ortlund, Stuart C. Sealfon, Michael P. Snyder, Martin J. Walsh

**Clinical Sites:** Cheehoon Ahn, Alicia Belangee, Bryan C. Bergman, Daniel H. Bessesen, Gerard A. Boyd, Anna R. Brandt, Nicholas T. Broskey, Toby L. Chambers, Clarisa Chavez Martinez, Maria Chikina, Alex Claiborne, Zachary S. Clayton, Paul M. Coen, Katherine A. Collins-Bennett, Tiffany M. Cortes, Gary R. Cutter, Matthew Douglass, Daniel E. Forman, Will A. Fountain, Aaron H. Gouw, Kevin J. Gries, Fadia Haddad, Joseph A. Houmard, Kim M. Huffman, Ryan P. Hughes, John M. Jakicic, Catherine M. Jankowski, Neil M. Johannsen, Johanna L. Johnson, Erin E. Kershaw, Dillon J. Kuszmaul, Bridget Lester, Colleen E. Lynch, Edward L. Melanson, Cristhian Montenegro, Kerrie L. Moreau, Masatoshi Naruse, Bradley C Nindl, Tuomo Rankinen, Ulrika Raue, Ethan Robbins, Kaitlyn R. Rogers, Renee J. Rogers, Irene E. Schauer, Robert S. Schwartz, Chad M. Skiles, Lauren M. Sparks, Maja Stefanovic-Racic, Andrew M. Stroh, Kristen J. Sutton, Anna Thalacker-Mercer, Todd A. Trappe, Caroline S. Vincenty, Elena Volpi, Katie L. Whytock, Gilhyeon Yoon, Thomas W. Buford, Dan M. Cooper, Sara E. Espinoza, Bret H. Goodpaster, Wendy M. Kohrt, William E. Kraus, Nicolas Musi, Shlomit Radom-Aizik, Blake B. Rasmussen, Eric Ravussin, Scott Trappe

**MoTrPAC Study Group Acknowledgements:** Nicole Adams, Abdalla Ahmed, Andrea Anderson, Carter Asef, Arianne Aslamy, Marcas M. Bamman, Jerry Barnes, Susan Barr, Kelsey Belski, Will Bennett, Amanda Boyce, Brandon Bukas, Emily Carifi, Chih-Yu Chen, Haiying Chen, Shyh-Huei Chen, Samuel Cohen, Audrey Collins, Gavin Connolly, Elaine Cornell, Julia Dauberger, Carola Ekelund, Shannon S. Emilson, Jerome Fleg, Nicole Gagne, Mary-Catherine George, Ellie Gibbons, Jillian Gillespie, Aditi Goyal, Bruce Graham, Xueyun Gulbin, Jere Hamilton, Leora Henkin, Andrew Hepler, Lidija Ivic, Ronald Jackson, Andrew Jones, Lyndon Joseph, Leslie Kelly, Gary Lee, Adrian Loubriel, Ching-ju Lu, Kristal M. Maner-Smith, Ryan Martin, Padma Maruvada, Alyssa Mathews, Curtis McGinity, Lucas Medsker, Kiril Minchev, Samuel G. Moore, Michael Muehlbauer, Anne Newman, John Nichols, Concepcion R. Nierras, George Papanicolaou, Lorrie Penry, June Pierce, Megan Reaves, Eric W. Reynolds, Teresa N. Richardson, Jeremy Rogers, Scott Rushing, Santiago Saldana, Rohan Shah, Samiya M. Shimly, Cris Slentz, Deanna Spaw, Debbie Steinberg, Suchitra Sudarshan, Alyssa Sudnick, Jennifer W. Talton, Christy Tebsherani, Nevyana Todorova, Mark Viggars, Jennifer Walker, Michael P. Walkup, Anthony Weakland, Gary Weaver, Christopher Webb, Sawyer Welden, John P. Williams, Marilyn Williams, Leslie Willis, Yi Zhang, Frank Booth, Karyn A. Esser, Laurie Goodyear, Andrea Hevener, Ian Lanza, Jun Li, K Sreekumaran Nair

