## Supplemental Figures for "Exercise modulation of the alternative splicing landscape in human tissues"

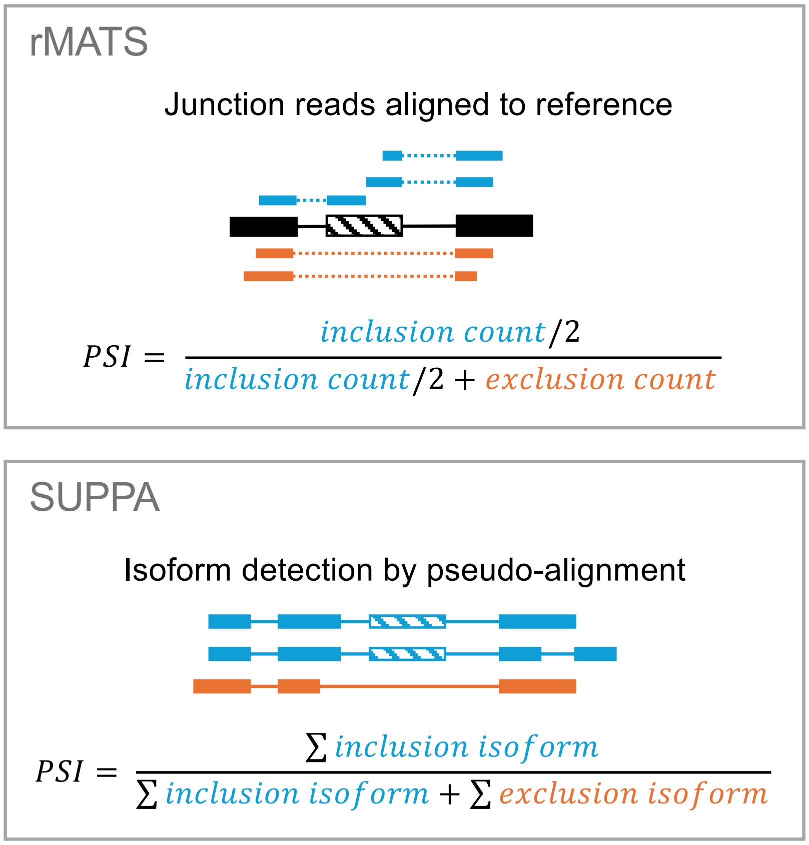


Supplementary Figure 1

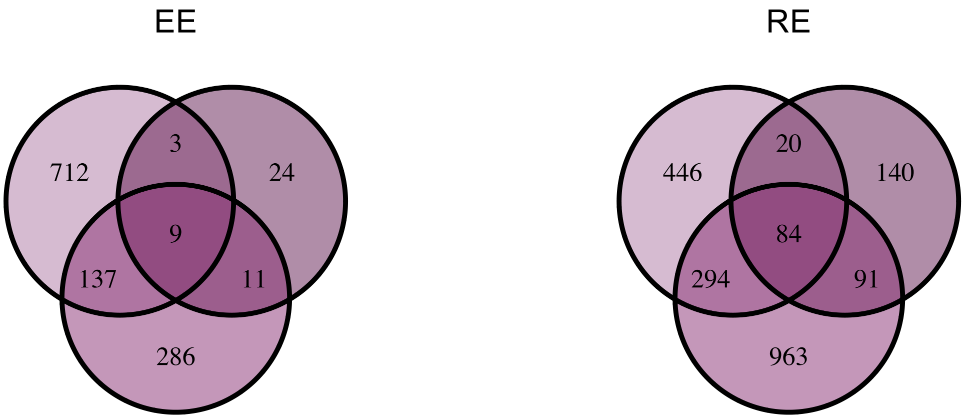


Supplementary Figure 2

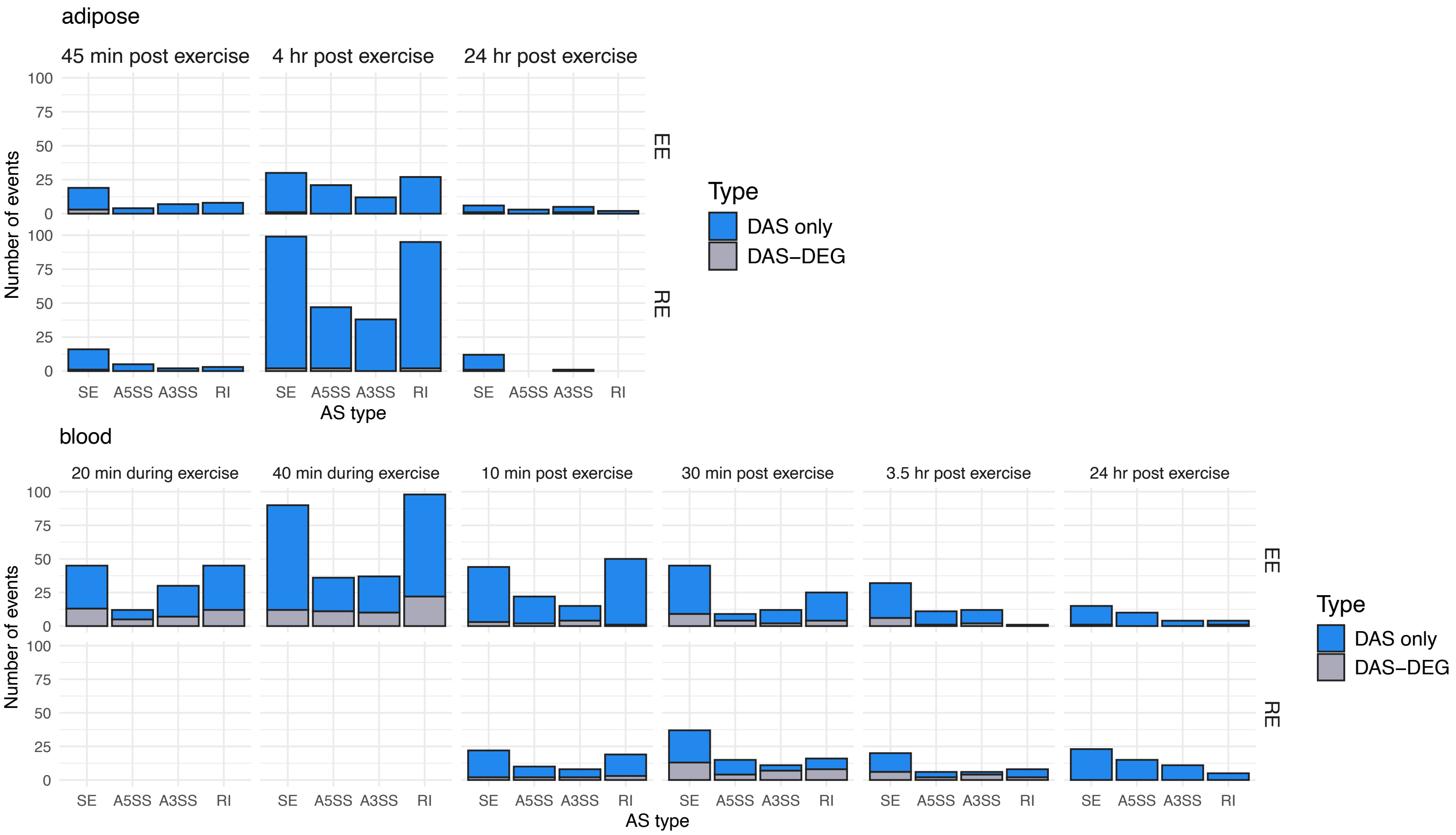


### Supplementary Figure 3

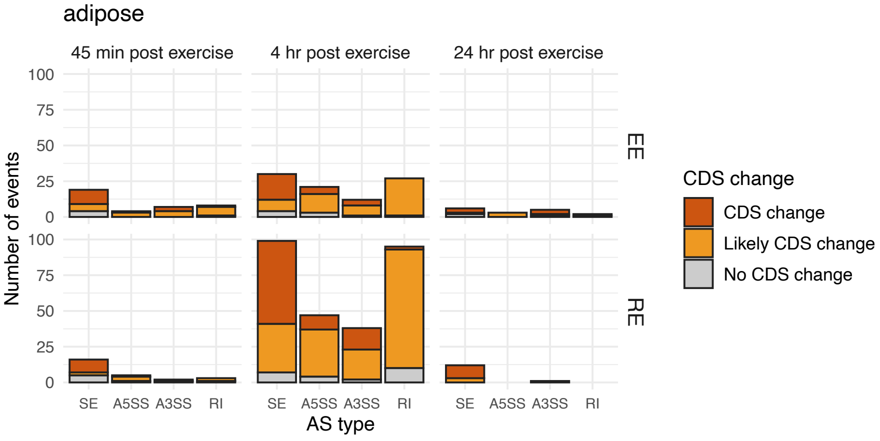


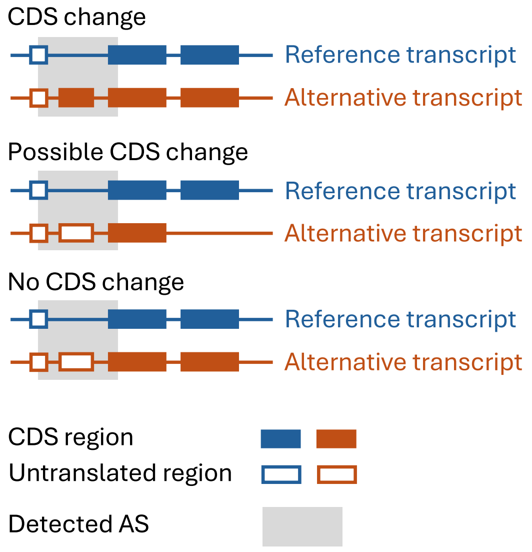


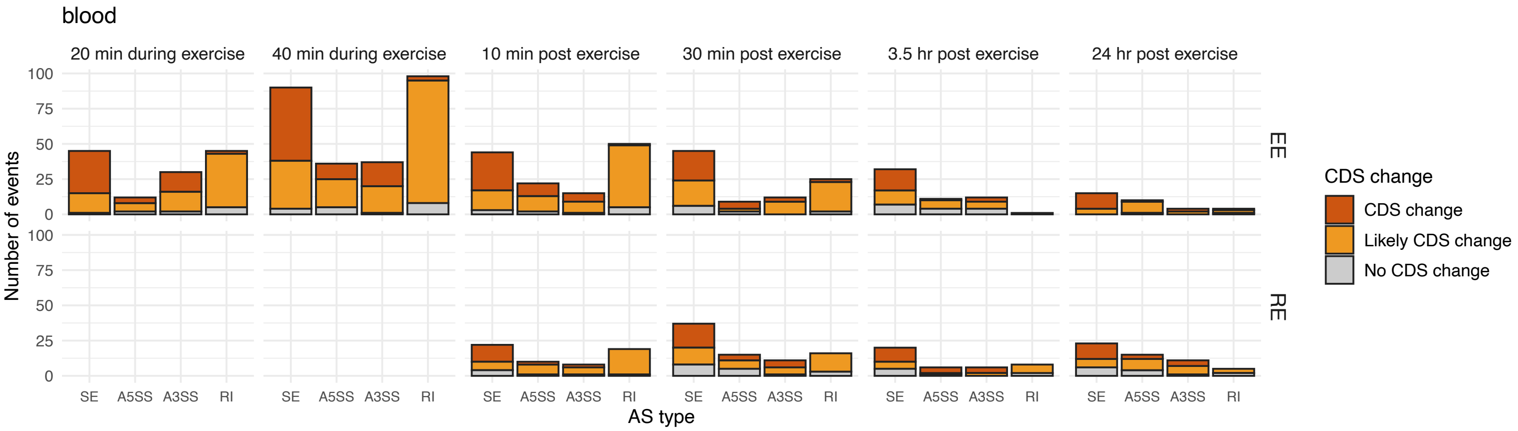


### Supplementary Figure 4

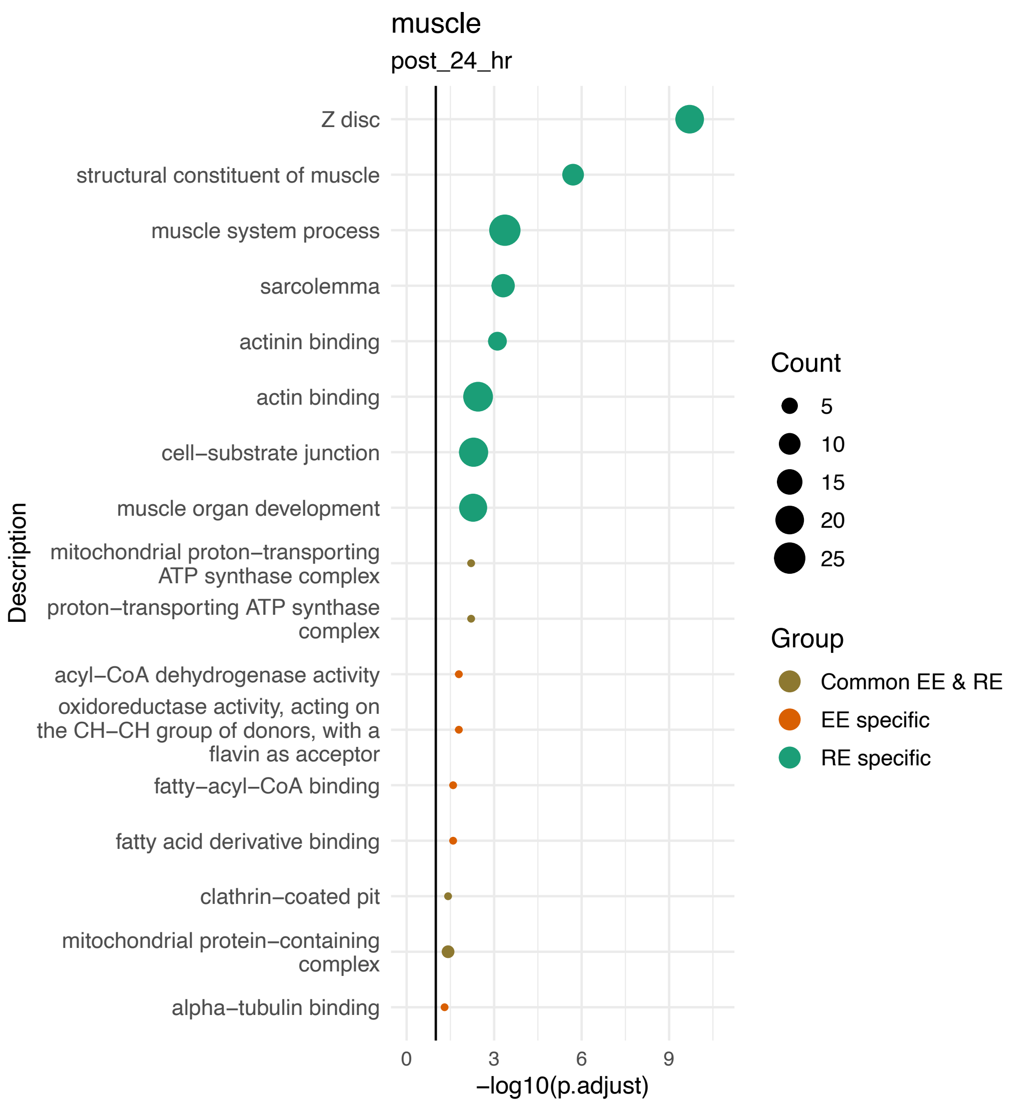


Supplementary Figure 5

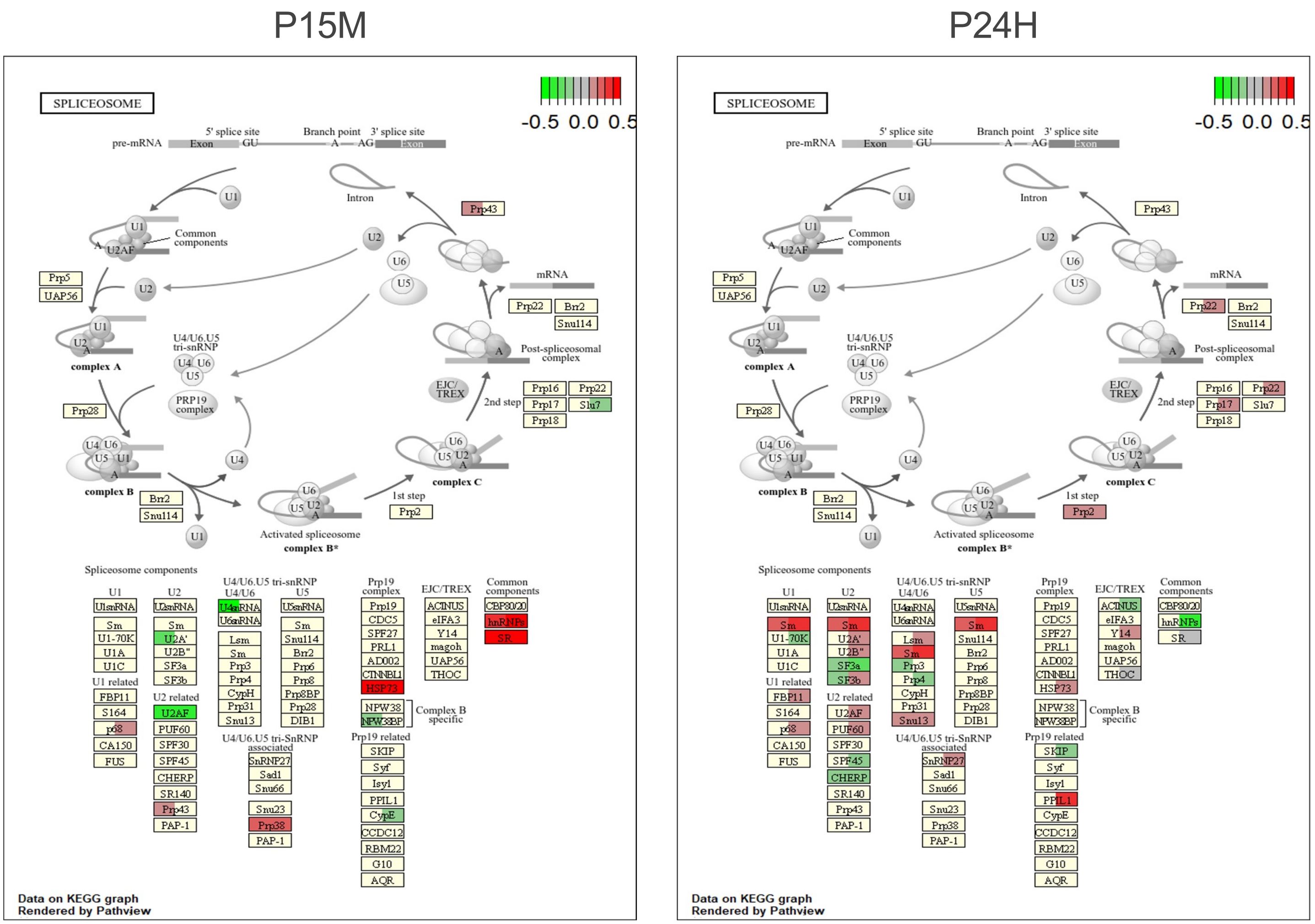


Supplementary Figure 6

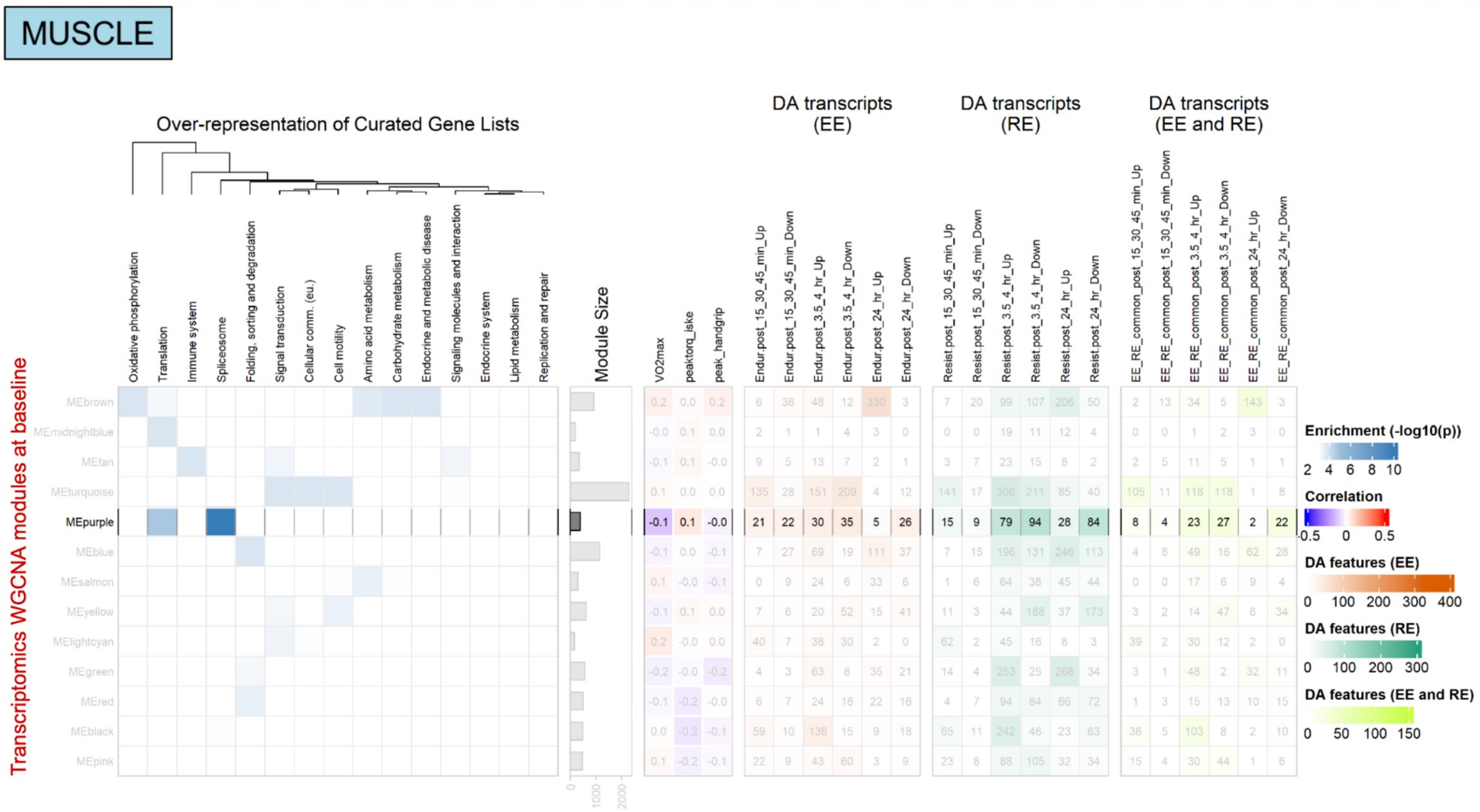


Supplementary Figure 7

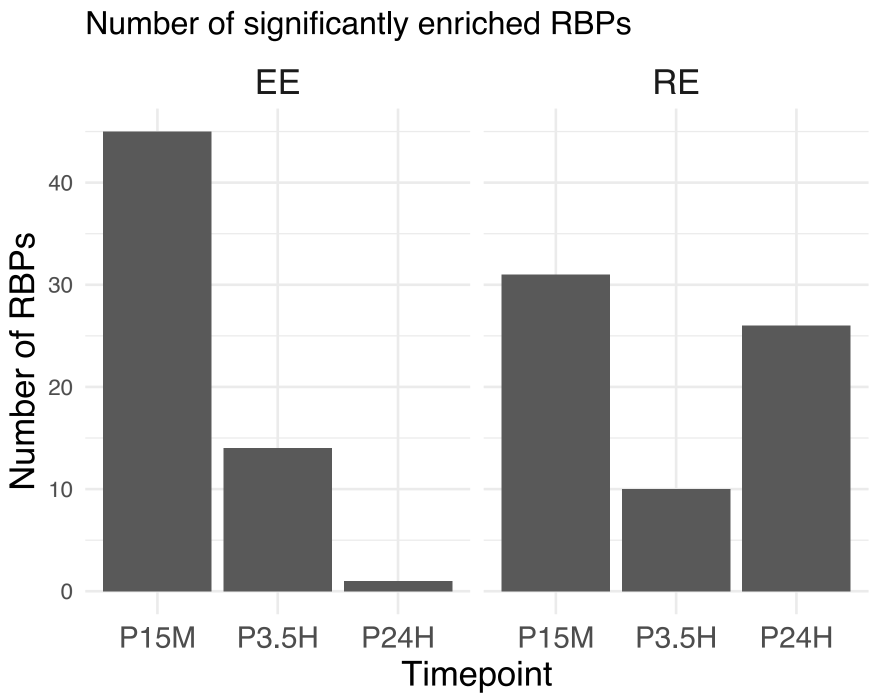


Supplementary Figure 8

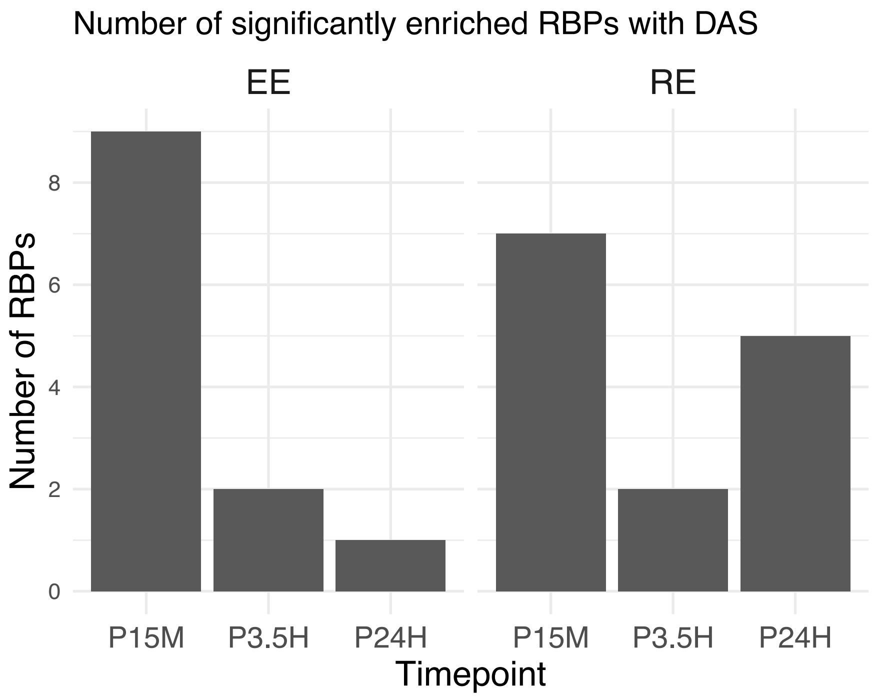


Supplementary Figure 9
